# Pan-cancer analysis of whole genomes reveals driver rearrangements promoted by LINE-1 retrotransposition in human tumours

**DOI:** 10.1101/179705

**Authors:** Bernardo Rodriguez-Martin, Eva G. Alvarez, Adrian Baez-Ortega, Jorge Zamora, Fran Supek, Jonas Demeulemeester, Martin Santamarina, Young Seok Ju, Javier Temes, Daniel Garcia-Souto, Harald Detering, Yilong Li, Jorge Rodriguez-Castro, Ana Dueso-Barroso, Alicia L. Bruzos, Stefan C. Dentro, Miguel G. Blanco, Gianmarco Contino, Daniel Ardeljan, Marta Tojo, Nicola D. Roberts, Sonia Zumalave, Paul A. W. Edwards, Joachim Weischenfeldt, Montserrat Puiggros, Zechen Chong, Ken Chen, Eunjung Alice Lee, Jeremiah A. Wala, Keiran Raine, Adam Butler, Sebastian M. Waszak, Fabio C. P. Navarro, Steven E. Schumacher, Jean Monlong, Francesco Maura, Niccolo Bolli, Guillaume Bourque, Mark Gerstein, Peter J. Park, David Wedge, Rameen Beroukhim, David Torrents, Jan O. Korbel, Inigo Martincorena, Rebecca C. Fitzgerald, Peter Van Loo, Haig H. Kazazian, Kathleen H. Burns, Peter J. Campbell, Jose M. C. Tubio, on behalf of the PCAWG Structural Variation Working Group, the ICGC/TCGA Pan-Cancer Analysis of Whole Genomes Network

**Affiliations:** Mobile Genomes and Disease, Molecular Medicine and Chronic diseases Centre (CIMUS), Universidade de Santiago de Compostela, Santiago de Compostela 15706, Spain; Department of Zoology, Genetics and Physical Anthropology, Universidade de Santiago de Compostela, Santiago de Compostela 15706, Spain; The Biomedical Research Centre (CINBIO), University of Vigo, Vigo 36310, Spain; Transmissible Cancer Group, Department of Veterinary Medicine, University of Cambridge, Cambridge CB3 0ES, UK; Genome Data Science, Institute for Research in Biomedicine IRBB, The Barcelona Institute of Science and Technology (BIST), Barcelona 08028, Spain; The Francis Crick Institute, London, UK; Department of Human Genetics, University of Leuven, Leuven, Belgium; Graduate School of Medical Science and Engineering, Korea Advanced Institute of Science and Technology, Daejeon 34141, Korea; Evolutionary Genomics, The Biomedical Research Centre - CINBIO, University of Vigo, 36310 Vigo, Spain; Cancer Ageing and Somatic Mutation Programme, Wellcome Trust Sanger Institute, Hinxton, Cambridge CB101SA, UK; Barcelona Supercomputing Center (BSC-CNS), Barcelona, Spain; Faculty of Science and Technology. University of Vic - Central University of Catalonia (UVic-UCC), Vic, Spain; Experimental Cancer Genetics, Wellcome Trust Sanger Institute, Hinxton, Cambridge CB10 1SA. UK; Oxford Big Data Institute, University of Oxford, Oxford, UK; Molecular Medicine and Chronic diseases Centre (CIMUS) – IDIS, University of Santiago de Compostela, Santiago de Compostela 15706, Spain; Departamento de Bioquímica e Bioloxía Molecular, CIMUS, Universidade de Santiago de Compostela, 15706 Santiago de Compostela, Spain; Medical Research Council (MRC) Cancer Unit, University of Cambridge, Cambridge, UK; Institute for Genetic Medicine, Johns Hopkins University School of Medicine - Baltimore, MD USA; Department of Pathology, University of Cambridge, Cambridge, UK; Cancer Research UK Cambridge Institute, Cambridge, UK; Biotech Research & Innovation Centre (BRIC); Finsen Laboratory, Rigshospitalet, Copenhagen, Denmark; European Molecular Biology Laboratory (EMBL), Genome Biology Unit, 69117 Heidelberg, Germany; Department of Bioinformatics and Computational Biology, The University of Texas MD Anderson Cancer Center, Houston, Texas, USA; Division of Genetics and Genomics, Boston Children’s Hospital, Harvard Medical School, Boston, MA, USA; The Broad Institute of Harvard and MIT, Cambridge, MA 02142, USA; Department of Cancer Biology, Dana-Farber Cancer Institute, Boston, MA 02215, USA; Program in Computational Biology and Bioinformatics, Yale University, Bass 432, 266 Whitney Avenue, New Haven, CT; Department of Molecular Biophysics and Biochemistry, Yale University, Bass 432, 266 Whitney Avenue, New Haven, CT; Department of Human Genetics, McGill University, Montreal, H3A 1B1, Canada; Department of Oncology and Onco-Hematology, University of Milan, Milan, Itlay; Department of Medical Oncology and Hematology, Fondazione IRCCS Istituto Nazionale dei Tumori, Milan, Italy; Oxford NIHR Biomedical Research Centre, Oxford, UK; Institucio Catalana de Recerca i Estudis Avançats (ICREA), Barcelona, Spain; Department of Pathology, Johns Hopkins University School of Medicine - Baltimore, MD USA; Department of Haematology, University of Cambridge, Cambridge CB2 2XY, UK

## Abstract

About half of all cancers have somatic integrations of retrotransposons. To characterize their role in oncogenesis, we analyzed the patterns and mechanisms of somatic retrotransposition in 2,954 cancer genomes from 37 histological cancer subtypes. We identified 19,166 somatically acquired retrotransposition events, affecting 35% of samples, and spanning a range of event types. L1 insertions emerged as the first most frequent type of somatic structural variation in esophageal adenocarcinoma, and the second most frequent in head-and-neck and colorectal cancers. Aberrant L1 integrations can delete megabase-scale regions of a chromosome, sometimes removing tumour suppressor genes, as well as inducing complex translocations and large-scale duplications. Somatic retrotranspositions can also initiate breakage-fusion-bridge cycles, leading to high-level amplification of oncogenes. These observations illuminate a relevant role of L1 retrotransposition in remodeling the cancer genome, with potential implications in the development of human tumours.

---

Long interspersed nuclear element (LINE)-1 (L1) retrotransposons are widespread repetitive elements in the human genome, representing 17% of the entire DNA content^1,2^. Using a combination of cellular enzymes and self-encoded proteins with endonuclease and reverse transcriptase activity, L1 elements copy and insert themselves at new genomic sites, a process called retrotransposition. Most of the ∼500,000 L1 copies in the human reference genome are truncated, inactive elements not able to retrotranspose. A small subset of them, maybe 100-150 L1 loci, remain active in the average human genome, acting as source elements, of which a small number are highly active copies termed hot-L1s^3-5^. These L1 source elements are usually transcriptionally repressed, but epigenetic changes occurring in tumours may promote their expression and allow them to retrotranspose^6,7^. Somatic L1 retrotransposition most often introduces a new copy of the 3’ end of the L1 sequence, and can also mobilize unique DNA sequences located immediately downstream of the source element, a process called 3’ transduction^7-9^. L1 retrotransposons can also promote the somatic trans-mobilization of Alu, SVA and processed pseudogenes, which are copies of messenger RNAs that have been reverse transcribed into DNA and inserted into the genome using the machinery of active L1 elements^10-12^.

Approximately 50% of human tumours have somatic retrotransposition of L1 elements^7,13-15^. Previous analyses indicate that although a fraction of somatically acquired L1 insertions in cancer may influence gene function, the majority of retrotransposon integrations in a single tumour represent passenger mutations with little or no effect on cancer development^7,13^. Nonetheless, L1 insertions are capable of promoting other types of genomic structural alterations in the germline and somatically, apart from canonical L1 insertion events^16-18^, which remain largely unexplored in human cancer^19,20^.

To further understand the roles of retrotransposons in cancer, we developed novel strategies to analyze the patterns and mechanisms of somatic retrotransposition in 2,954 cancer genomes from 37 histological cancer subtypes within the framework of the Pan-Cancer Analysis of Whole Genomes (PCAWG) project ^21^, many of which have not been previously evaluated for retrotransposition. This work illuminates novel, hidden patterns and mutational mechanisms of structural variation in human cancers mediated by L1 retrotransposition. We find that aberrant integration of L1 retrotransposons has a relevant role in remodeling cancer genome architecture in some human tumours, mainly by promoting megabase-scale deletions that, occasionally, generate genomic consequences that may promote cancer development through the removal of tumour suppressor genes, such as *CDKN2A*, or triggering amplification of oncogenes, such as *CCND1*.

## RESULTS

### The landscape of somatic retrotransposition in the largest cancer whole-genomes dataset

We ran our bioinformatic pipelines to explore somatic retrotransposition on whole genome sequencing data from 2,954 tumours and their matched normal pairs, across 37 cancer types (**Supplementary Fig. 1** and **Supplementary Table 1**). The analysis retrieved a total of 19,166 somatically acquired retrotranspositions that were classified into six categories (Fig. 1a and **Supplementary Table 2**). With 98% (18,739/19,166) of the events, L1 integrations (14,967 Solo-L1, 3,669 L1-transductions, and 103 L1-mediated rearrangements – mainly deletions) overwhelmingly dominate the landscape of somatic retrotransposition in the PCAWG dataset. The vast majority of these events (97.5%) belong to the youngest L1 subfamilies Ta-1 and Ta-0 (**Supplementary Fig. 2**). Elements of the lineages Alu and SVA, with 130 and 23 somatic copies, represent minor categories in the somatic retrotransposition landscape (Fig. 1a and **Supplementary Table 2**). Despite the fact that we identified novel germline polymorphic integrations of endogenous retroviruses (ERVs) in the matched-normal samples cohort ^22^, no somatic ERV events are detected in any of the PCAWG tumours. We find that trans-mobilization of processed pseudogenes, with 274 events, is also a rare class of retrotransposition in the PCAWG dataset. Nonetheless, genomic landscapes in the cohort reveal that pseudogenes may have a particularly strong representation among retrotransposition events. For example, one relevant pancreatic cancer, SA533710, harbors ∼26% (70/274) of all processed pseudogenes mobilized somatically in the PCAWG cohort (Fig. 1b).

**Figure 1.**
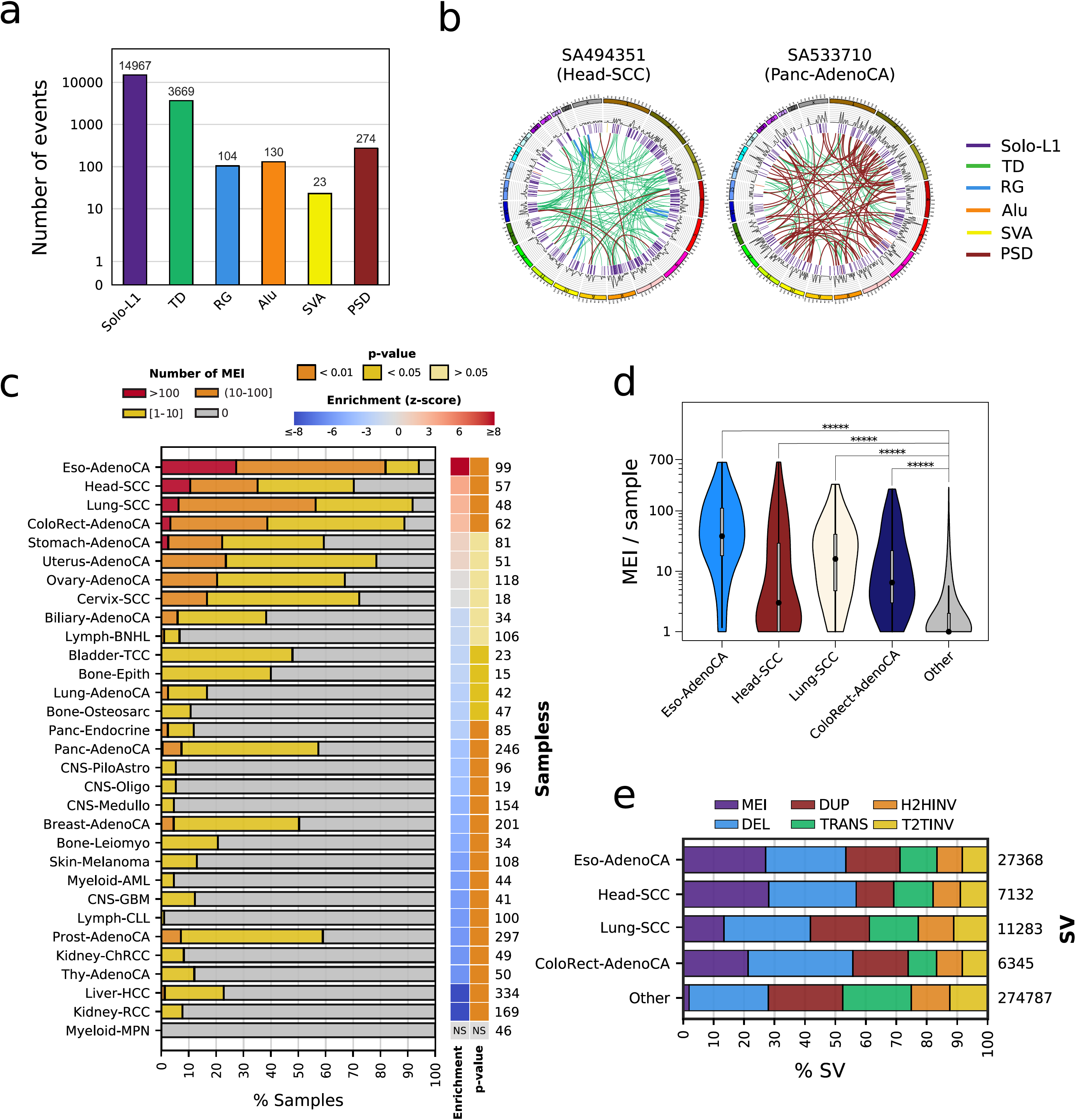
Landscape of somatic retrotransposition across human cancers. (a) Histogram showing the distribution of the number of somatic mobile element insertions (MEI) observed in 2,954 cancer genomes across six retrotransposition categories: Solo-L1, L1-mediated transductions (TD), L1-mediated rearrangements (RG), Alu, SVA, Pseudogenes (PSD). (b) Circos plots showing the genomic landscape of somatic retrotransposition in two representative tumours with high L1 activity. Left, a head-and-neck squamous carninoma, SA494351, bears 638 somatic insertion events, being the PCAWG sample with the highest retrotransposition rate. Right, a single pancreatic adenocarcinoma sample, SA533710, harbours ∼26% of all processed pseudogenes identified in the PCAWG cohort. Chromosome ideograms are shown around the outer ring with individual rearrangements shown as arcs, each coloured according to the type of rearrangement. (c) For 31 well-represented (i.e., with a minimum sample size of 15) cancer types included in the PCAWG dataset, it is represented the proportion of tumour samples with more than 100 somatic retrotransposon integrations (red), from 10 to 100 (orange), between 1 and 10 (yellow), and zero (grey). Retrotransposition enrichment or depletion for each tumour type together with level of significance (Zero-inflated negative binomial regression) are shown. “NS” stands for non-significant. The number of samples analyzed from each tumour type is shown on the right side of the panel. (d) Distribution of retrotransposition events per sample across cancer types. Only the four tumour types significantly enriched in retrotransposition are shown, the remaining are grouped into the category ‘Other’. Distribution between each cancer and ‘Other’ is compared though Mann–Whitney U; p-values lower than 0.0001 are indicated with five asterisks. Y-axis is presented in a logarithmic scale. An extended version of this panel is shown in **Supplementaty Fig. 5**. (e) For the same four tumour types from panel d, it is shown the fraction of structural variants (SVs) belonging to the six different SV classes catalogued in PCAWG ^45^. Mobile element insertions (MEI), deletions (DEL), duplications (DUP), translocations (TRANS), head-2-head inversions (H2HINV) and tail-2-tail inversions (T2TINV). The total number of variants observed per cancer type is indicated on the right side of the panel.

The core pipeline, TraFiC-mem (**Supplementary Fig. 3**), employed to explore all classes of somatic retrotransposition in the PCAWG dataset, was validated by single-molecule whole-genome sequencing data analysis from one cancer cell-line with high retrotransposition rate and its matched normal sample, confirming the somatic acquisition of 295 out of 308 retrotransposition events (false discovery rate <5%, **Supplementary Fig. 4a-b**). To further evaluate TraFiC-mem, we reanalyzed a mock cancer genome into which we had previously ^7^ seeded somatic retrotransposition events at different levels of tumour clonality, and then simulated sequencing reads to the average level of coverage of the PCAWG dataset. The results confirmed a high precision (>99%) of TraFiC-mem, and a recall ranging from 90 to 94% for tumour clonalities from 25 to 100%, respectively (**Supplementary Fig. 4c-e**).

We observe a dramatic variation of the retrotransposition rate across PCAWG tumour types (Fig. 1c and **Supplementary Table 3**). Overall, 35% (1,046/2,954) of all cancer genomes have at least one retrotransposition event. However, esophageal adenocarcinoma, head-and-neck squamous carcinoma, lung squamous carcinoma, and colorectal adenocarcinoma, are significantly enriched in somatic retrotranspositions (Mann–Whitney U test, P < 0.05; Fig. 1c-d and **Supplementary Fig. 5**). These four tumour types alone account for 70% (13,373/19,166) of all somatic events in the PCAWG dataset, although they represent just 9% (266/2,954) of the samples. This is particularly noticeable in esophageal adenocarcinoma, where 27% (27/99) of the samples show more than 100 separate somatic retrotranspositions (Fig. 1c), making L1 insertions the most frequent type of structural variation in esophageal adenocarcinoma (Fig. 1e). Furthermore, retrotranspositions are the second most frequent type of structural variants in head-and-neck squamous and colorrectal adenocarcinomas (Fig. 1e). In order to gain insights into the genetic causes that make some cancers more prone to retrotransposition than others, we looked for associations between retrotransposition and driver mutations in cancer-related genes. This analysis revealed an increased L1 retrotransposition rate in tumours with *TP53* mutations (Mann–Whitney *U* test, *P* < 0.05; **Supplementary Fig. 6**), and supports previous analysis that suggested *TP53* functions to restrain mobile elements ^23,24^. We also observe a widespread correlation between L1 retrotransposition and other types of structural variation (Spearman’s ρ = 0.44, P < 0.01, **Supplementary Fig 7**), a finding most likely consequence of a confounding effect of *TP53*-mutated genotypes, given by the fact that *TP53* mutants also exhibit an increased number of structural variants (Mann–Whitney *U* test, *P* < 0.05; **Supplementary Fig. 6**).

We identify 43% (7,979/18,636) somatic retrotranspositions of L1 inserted within gene regions including promoters, of which 66 events hit cancer genes. The analysis of expression levels in samples with available transcriptome, revealed four genes with L1 retrotranspositions in the proximity of promoter regions showing significant over-expression when compared to the expression in the remaining samples from the same tumour type (Student’s t-test, q < 0.10; **Supplementary Fig. 8a-c**). This includes one head-and-neck tumour, SA494343, with a somatic L1 element integrated into the promoter region of the *ABL2* oncogene. The structural analysis of RNA-seq data identified instances in which portions of a somatic retrotransposition within a gene exonize, being incorporated into the transcript sequence of the affected gene, a process that sometimes involved cancer genes (**Supplementary Fig. 8d**). In addition, we analyzed the potential of processed pseudogenes to generate functional consequences in cancer^11^. We found evidence for aberrant fusion transcripts arising from inclusion of 14 processed pseudogenes in the target host gene, and expression of 3 processed pseudogenes landing in intergenic regions (**Supplementary Fig. 8e**).

### Dissecting the genomic features that influence the landscape of L1 retrotranspositions in cancer

The genome-wide analysis of the distribution of somatic L1 insertions across the cancer genome, revealed a dramatic variation of L1 retrotransposition rate (Fig 2a and **Supplementary Table 4**). To understand the reasons of such variation, we studied the association of L1 event rates with various genomic features. We first wondered if the distribution of somatic L1s across the cancer genome could be determined by the occurrence of L1-endonuclease target site motifs. We used a statistical approach based on negative binomial regression to deconvolute the influence of multiple overlapping genomic variables^25^, which showed that close matches to the motif have a 244-fold increased L1 rate, compared to non-matched motifs (Fig. 2b and **Supplementary Fig. 9a**). Adjusting for this effect, we find a strong association with DNA replication time, with the latest-replicating quarter of the genome being 8.9-fold enriched in L1 events (95% CI: 8.25-9.71) compared to the earliest-replicating quarter (Fig. 2b-c and **Supplementary Fig. 9b**). Recent work ^26^ has shown that L1 retrotransposition has a strong cell cycle bias, preferentially occuring in the S-phase. Our results are in agreement with these findings, and suggest that L1 retrotransposition peaks in the later stages of the nuclear DNA synthesis.

**Figure 2.**
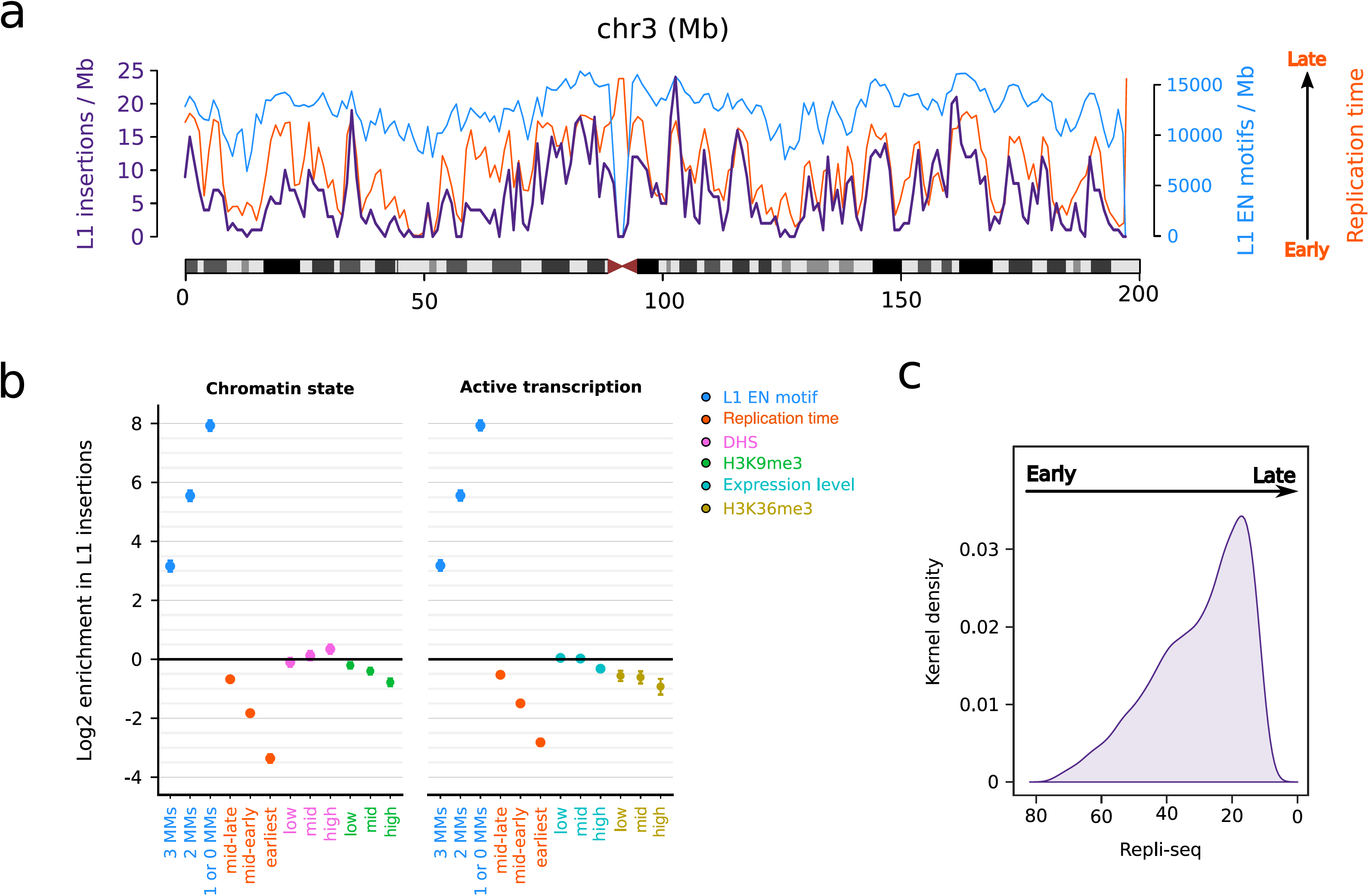
Distribution of L1 somatic insertions across the cancer genome and its association with genome organization features. We performed a genome-wide analysis of the distribution of 15,906 somatic L1 insertions, which include solo-L1 and L1 transductions with a 3’-poly(A) breakpoint characterized to base-pair resolution. (a) This analysis revealed a dramatic variation of L1 retrotransposition rate across the cancer genome. Here, L1 insertion rate (purple) is shown together with L1 endonuclease (EN) motif density (blue), and replication timing (orange). The data is represented per 1 Mb-window. For illustrative purposes, only chromosome 3 is shown; the full data is presented in **Supplementary Table 4**. (b) Association between L1 insertion rate and multiple predictor variables at single-nucleotide resolution. All enrichment scores shown are adjusted for multiple covariates and compare the L1 insertion rate in bins 1-3 for a particular genomic feature (L1 EN motif, replication timing, open chromatin, histone marks and expression level) versus bin 0 of the same feature, which therefore always has log enrichment=0 by definition and is not shown. MMs stands for the number of mismatches with respect the consensus L1 EN motif (see Methods). Heterochromatic regions and transcription elongation are defined based on H3K9me3 and H3K36me3 histone marks. Accessible chromatin is measured via DNase hypersensitivity (DHS). (c) L1 insertion density, using Kernel density estimate (KDE), along replication timing spectrum. DNA replication timing is expressed on a scale from 80 (early) to 0 (late).

Then, we examined L1 rates in open chromatin measured via DNase hypersensitivity (DHS) and, conversely, in closed-heterochromatic regions with the H3K9me3 histone mark ^27^. When adjusting for the confounding effects of L1 motif content and replication time ^25^, we find that somatic L1 events are enriched in open chromatin (1.27-fold in the top DHS bin, 95% CI 1.14-1.41; Fig. 2b), and depleted in heterochromatin (1.72-fold, 95% CI 1.57-1.99; Fig. 2b). This finding differs from previous analysis that suggested L1 insertions favored heterochomatin ^7^, a discrepancy we believe to be due to the confounding effect between heterochromatin and late-replicating DNA regions, which was not addressed in previous analyses. We also find negative association of L1 rates with active transcription chromatin features, characterized by fewer L1 events at active promoters (1.63-fold, **Supplementary Fig. 9c**), a slightly but significantly reduced L1 rates in highly expressed genes (1.25-fold lower, 95% CI: 1.16-1.34; Fig. 2d) and a further depletion at H3K36me3 (1.90-fold reduction in the highest tertile; 95% CI: 1.59-2.29; Fig. 2d), a mark of actively transcribed regions deposited in the body and the 3’ end of active genes ^27^. More details on these associations are shown in **Supplementary Fig. 9c-e**.

### L1 source elements’ contribution to Pan-Cancer retrotransposition burden

We used somatically mobilized L1-3’ transduction events to trace L1 activity to specific source elements^7^. This strategy revealed that 124 germline L1 loci in the human genome that are responsible for most of the genomic variation generated by retrotransposition in PCAWG ^7,22^ (**Supplementary Table 5**). Fifty-two of these loci represent novel, previously unreported source elements in human cancer ^22^. We analyzed the relative contribution of individual source elements to retrotransposition burden across cancer types, finding that retrotransposition is generally dominated by five hot-L1 source elements that alone give rise to half of all somatic transductions (Fig. 3a). This analysis revealed a dichotomous pattern of hot-L1 activity, with source elements we have termed Strombolian and Plinian, given their similarity to these two types of volcanos (Fig. 3b). Strombolian source elements are relatively indolent, producing small numbers of retrotranspositions in individual tumour samples, though they are often active and contribute significantly to overall retrotransposition in PCAWG. In contrast, Plinian elements are rarely active across tumours, but in these isolated cases, their activity is fulminant, causing large numbers of retrotranspositions.

**Figure 3.**
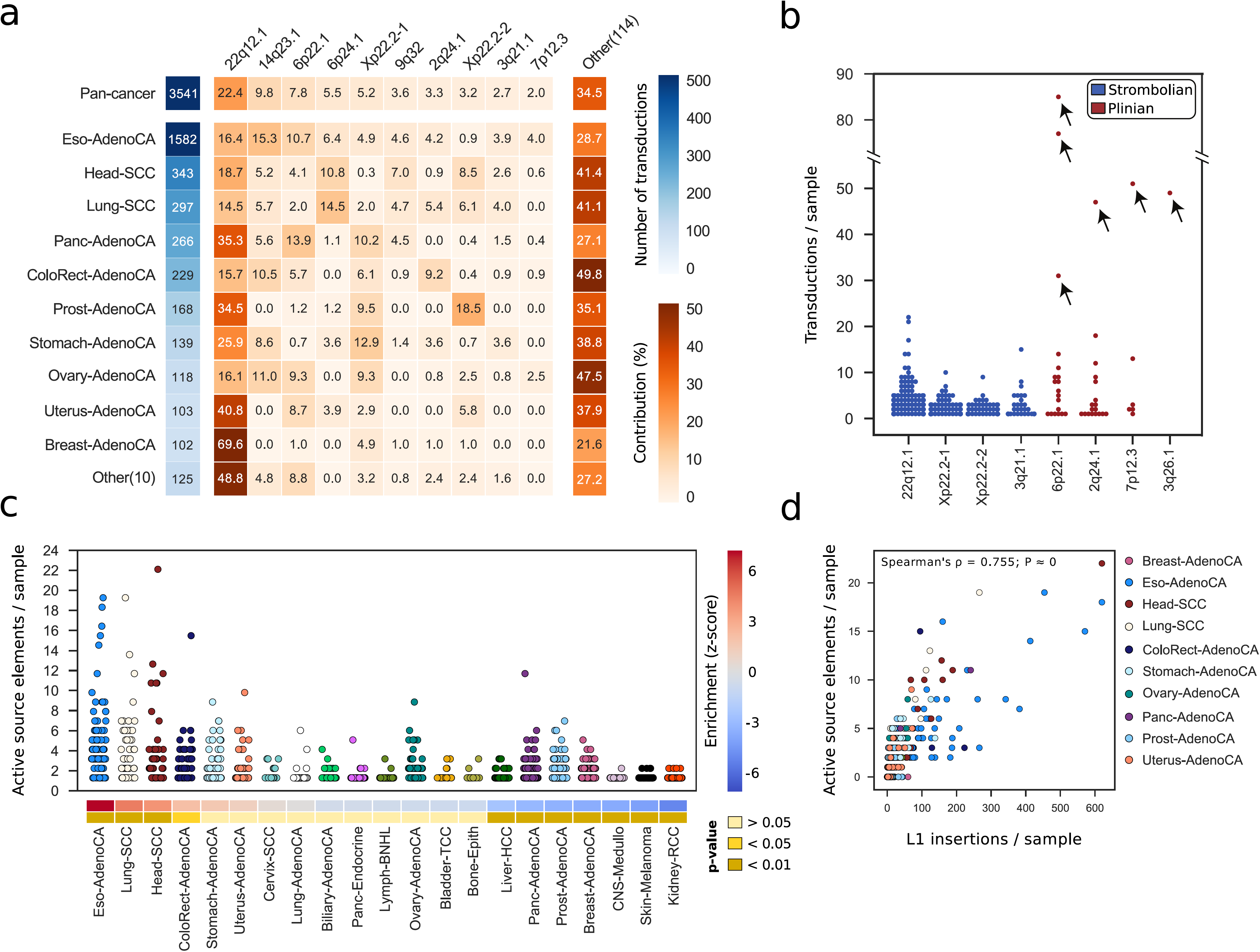
The dynamics of L1 source elements activity in human cancer. (a) We analyzed the contribution of 124 germline L1 source loci to somatic retrotransposition burden in PCAWG tumours. The total number of transductions identified for each cancer type is shown in a blue coloured scale. Contribution of each source element is defined as the proportion of the total number of transductions from each cancer type that is explained by each source loci. Only the top 10 contributing source elements are shown, while the remaining are grouped into the category ‘Other’. Additional information relative to these 10 loci and the remaining source elements can be obtained from **Supplementary Table 5**. (b) Two extreme patterns of hot-L1 activity, Strombolian and Plinian, were identified. Dots show the number of transductions promoted by each source element in a given tumour sample. Arrows indicate violent eruptions in particular samples (Plinian source elements). (c) Distribution of the number of active germline L1 source elements per sample, across cancer types with source element activity. Here, a source element is considered to be active in a given sample when it promotes at least one transduction in that particular sample. Number of active source elements enrichment or depletion for each tumour type together with level of significance (Zero-inflated negative binomial regression) are shown. (d) Correlation between the number of somatic L1 insertions and the number of active germline L1 source elements in PCAWG samples. Each dot represents a tumour sample and colours match cancer types.

At the individual tumour level, although we observe that the number of active source elements in a single cancer genome may vary from 1 to 22, typically only 1 to 3 loci are operative (Fig. 3c). There is a correlation of somatic retrotranspositions with number of active germline L1 source elements among PCAWG samples (Fig. 3d); this is likely one of the factors that explain why esophageal adenocarcinoma, lung and head-and-neck squamous carcinoma account for higher retrotransposition rates – in these three tumour types we also observed higher numbers of active germline L1 loci (Fig. 3c). Occasionally, somatic L1 integrations that retain their full length may also act as source for subsequent somatic retrotransposition events^7,28^, and may reach hot activity rates, leading them to dominate retrotransposition in a given tumour. For example, in a remarkable head-and-neck tumour, SA197656, we identify one somatic L1 integration at 4p16.1 that then triggers 18 transductions from its new site, with the next most active element being a germline L1 locus at 22q12.1 accounting for 15 transductions (**Supplementary Table 5**).

### Genomic deletions mediated by somatic L1 retrotransposition

In cancer genomes with high somatic L1 activity rates, we observed that some L1 retrotransposition events followed a distinctive pattern consisting of a single cluster of reads, associated with copy number loss, whose mates unequivocally identify one extreme of a somatic L1 integration with, apparently, no local, reciprocal cluster supporting the other extreme of the L1 insertion (Fig. 4a). Analysis of the associated copy number changes identified the missing L1 reciprocal cluster at the far end of the copy number loss, indicating that this pattern represents a deletion occurring in conjunction with the integration of an L1 retrotransposon (Fig. 4b). These rearrangements, called L1-mediated deletions, have been observed to occur somatically with engineered L1s in cultured human cells^16,17^ and naturally in the brain^18^, and are most likely the consequence of an aberrant mechanism of L1 integration.

**Figure 4.**
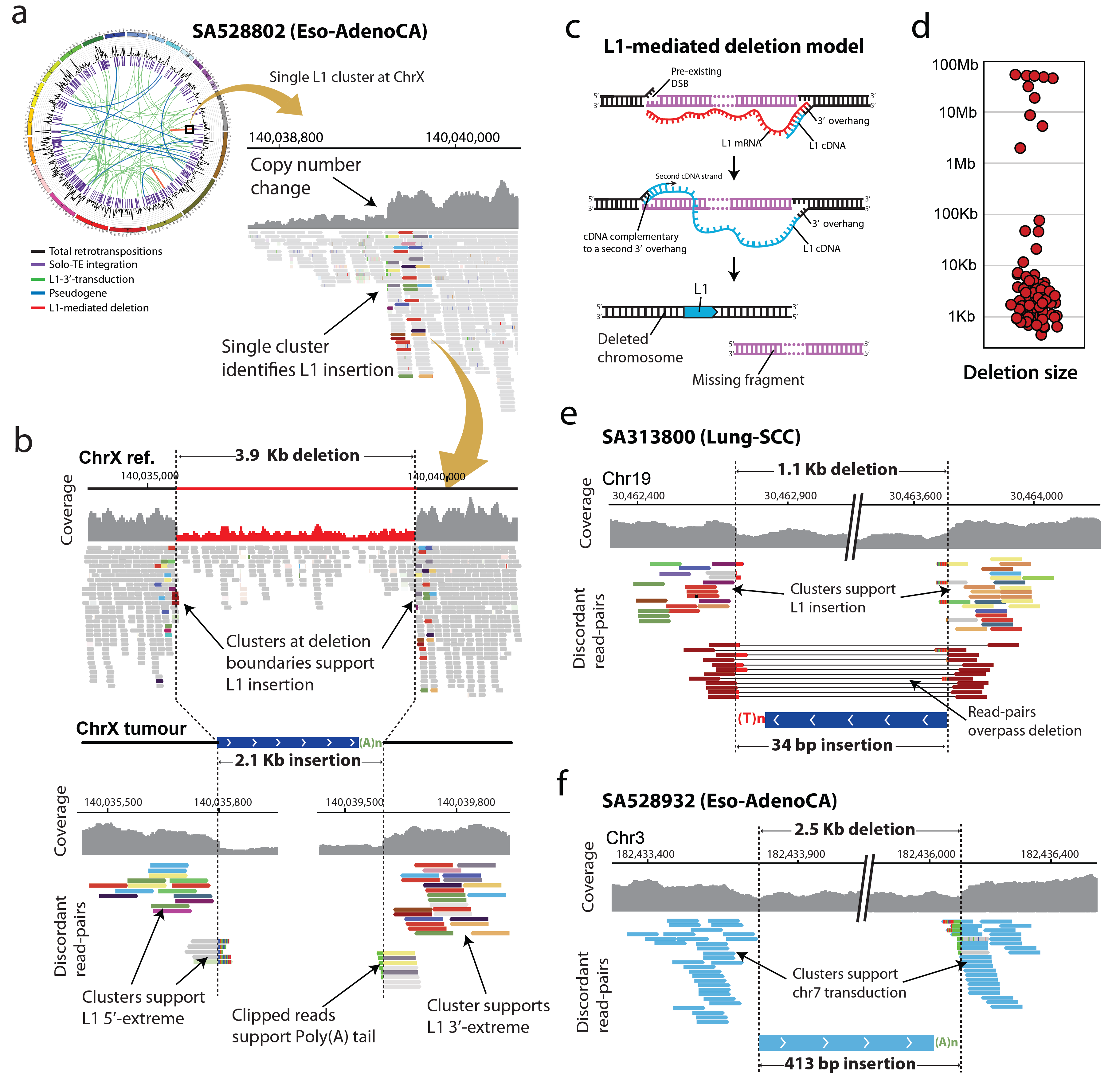
The hallmarks of somatic L1-mediated deletions revealed by copy number and paired-end mapping analysis. (a) In the retrotransposition analysis of SA528802, an esophageal tumour with high L1 somatic activity rates, we found one-single cluster of reads on chromosome X, which is associated with one breakpoint of a copy number loss, and whose mates unequivocally identify one extreme of a somatic L1 integration with, apparently, no local reciprocal cluster supporting the other extreme of the L1 insertion. This figure uses default read colours by Integrative Genomics Viewer (igv)^46^, where paired-end reads are coloured by the chromosome on which their mates can be found. Different colours for different reads from the same cluster indicate that mates are mapping a repetitive element. See **Supplementary Fig. 2a** for additional information on how to interpret sequencing data. (b) Analysis of the associated copy number change on chromosome X identifies the missing L1 reciprocal cluster far away, at the second breakpoint of the copy number loss, and reveals a 3.9 kb deletion occurring in conjunction with the integration of a 2.1 kb L1 somatic insertion. The sequencing data associated with this L1-mediated deletion shows two clusters of discordant read-pairs (i.e., pairs that have unexpected mapping distance) and clipped reads (i.e., reads that span one of the two insertion breakpoints) supporting both extremes of an L1 retrotransposon. (c) Model of L1-mediated deletion. The integration of an L1 mRNA typically starts with an L1-endonuclease cleavage promoting a 3’-overhang necessary for reverse transcription. Then, the cDNA (-) strand invades a second 3’-overhang from a pre-existing double-strand break upstream of the initial integration site. (d) Distribution of the sizes of 90 L1-mediated deletions identified in the PCAWG dataset. (e) In a Lung squamous carcinoma, SA313800, a 34 bp truncated L1 insertion promotes a 1.1 kb deletion at chromosome 19. Because the L1 insertion is so short, we also identify discordant read-pairs that span the L1 event and support the deletion. (f) In an esophageal adenocarcinoma, SA528932, the integration in chromosome 3 of a 413 bp orphan L1-transduction from chromosome 7 causes a 2.5 kb deletion, which is supported by two clusters of discordant read-pairs whose mates map onto the same region at chromosome 7.

We developed specific algorithms to systematically identify L1-mediated deletions, and applied this across all PCAWG tumours. This identified 90 somatic events matching the patterns described above causing deletions of different size, ranging from ∼0.5 kb to 53.4 Mb (Fig. 4d and **Supplementary Table 6**). The reconstruction of the sequence at the breakpoint junctions in each case supports the presence of an L1 element – or L1-transduction – sequence and its companion polyadenylate tract, indicative of passage through an RNA intermediate. No target site duplication is found, which is also the typical pattern for L1-mediated deletions^17^. One potential mechanism for these events is that a molecule of L1 cDNA pairs with a distant 3´-overhang from a preexisting double strand DNA break generated upstream of the initial integration site, and the DNA region between the break and the original target site is subsequently removed by aberrant repair^17^ (Fig. 4c). Indeed, in 75% (47/63) of L1-mediated deletions with a 5’-end breakpoint characterized to base-pair resolution, the analysis of the sequences at the junction revealed short (1-5 bp long, with median at 3 bp) microhomologies between the pre-integration site and the 5’ L1 sequence integrated right there (**Supplementary Table 6**). Furthermore, we find 14% (9/63) instances where short insertions (1-33 bp long, with median at 9 bp) are found at the 5’-breakpoint junction of the insertion. Both signatures are consistent with a non-homologous end-joining mechanism ^29^, or other type of microhomology-mediated repair, for the 5’-end attachment of the L1 cDNA to a 3´-overhang from a preexisting double-strand DNA break located upstream. L1-mediated deletions where microhomologies or insertions are not found may follow alternative models ^17,30-32^.

To confirm that these rearrangements are mediated by the integration of a single intervening retrotransposition event, we explored the PCAWG dataset for somatic L1-mediated deletions where the L1 sequences at both breakpoints of the deletion can be unequivocally assigned to the same L1 insertion. These include small deletions and associated L1 insertions shorter than the library size, allowing sequencing read-pairs to overlay the entire structure. For example, in a lung tumour, SA313800, we identified a deletion involving a 1 kb region at 19q12 with hallmarks of being generated by an L1 element (Fig. 4e). In this rearrangement we find two different types of discordant read-pairs at the deletion breakpoints: one cluster that supports the insertion of an L1 element, and a second that spans the L1 event and supports the deletion. Another type of L1-mediated deletion that can unequivocally be assigned to one-single L1 insertion event is represented by those deletions generated by the integration of orphan L1-transductions. These transductions represent fragments of unique DNA sequence located downstream of an active L1 locus, which are mobilized without the companion L1^7,15^. For example, in one oesophageal tumour, SA528932, we find a deletion of 2.5 kb on chromosome 3 mediated by the orphan transduction of sequence downstream of an L1 locus on chromosome 7 (Fig. 4f).

Due to the unavailability of PCAWG DNA specimens, we performed validation of 16 additional somatic L1-mediated deletions identified by TraFiC-mem in two head-and-neck cancer cell-lines with high retrotransposition rates, NCI-H2009 and NCI-H2087 ^7^. We carried out PCR followed by single-molecule sequencing of amplicons from these two tumour cell-lines and their matched normal samples, NCI-BL2009 and NCI-BL2087. The results confirmed that these rearrangements were somatically acquired, and that the insertion of an L1 event within the deletion boundaries was real (**Supplementary Fig. 10-11** and **Supplementary Table 7**). To further validate L1-mediated deletions, we performed Illumina whole-genome sequencing on the same two head-and-neck cancer cell-lines mentioned above, using mate-pair libraries with long insert sizes (3 kb and 10 kb) that would exceed the insertion event at the deletion boundaries. In these samples, our algorithms confirmed 16 events with the hallmarks of L1-mediated deletions, in which the mate-pair data confirmed a single L1-derived (i.e., solo-L1 or L1-transduction) retrotransposition as the cause of the copy number loss, and identified the sizes of the deletion and the associated insertion (**Supplementary Fig. 12**).

Analysis of L1 3’-extreme insertion breakpoint sequences from L1-mediated deletions found in the PCAWG dataset, revealed that 86% (74/90) of the L1 events causing deletions preferentially inserted into sequences that resemble L1-endonuclease consensus cleavage sites (e.g., 5’-TTTT/A-3’ and related sequences^33^) (**Supplementary Table 6)**. This confirms that L1 machinery, through a target-primed reverse transcription (TPRT) mechanism, is responsible for the integration of most of the L1 events causing neighboring DNA loss^33^. Interestingly, in 16% (14/90) of the events the endonucleotidic cleavage occurred at the phosphodiester bond between a T/G instead of at the standard T/A. In addition, we observed 8% (7/90) instances where the endonuclease motif is not found and the integrated element is truncated at both the 5′ and 3′ ends, suggesting that a small fraction of L1-associated deletions are the consequence of an L1-endonuclease-independent insertion mechanism^31-33^. Whatever mechanism of L1 integration operating here, taken together, these data indicate that the somatic integration of L1 elements induces the associated deletions.

### Megabase-size L1-mediated deletions cause loss of tumour suppressor genes

Most L1-mediated deletions ranged from a few hundred to thousands of base pairs, but occasionally deleted megabase regions of a chromosome (Fig. 4d and **Supplementary Table 6**). For example, in oesophageal tumour SA528901, we find a 45.5 Mb interstitial deletion involving the p31.3-p13.3 regions from chromosome 1 (Fig. 5a), where both breakpoints of the rearrangement show the hallmarks of a deletion mediated by integration of an L1 element. Here, the L1 element is 5’-truncated, which rendered a small L1 insertion, allowing a fraction of the sequencing read-pairs to span both breakpoints of the rearrangement. This unequivocally supports the model that the observed copy number change is indeed a deletion mediated by retrotransposition of an L1 element. Similarly, in a lung tumour, SA313800, we found an interstitial L1-mediated deletion with loss of 51.1 Mb from chromosome X including the centromere (Fig. 5b).

**Figure 5.**
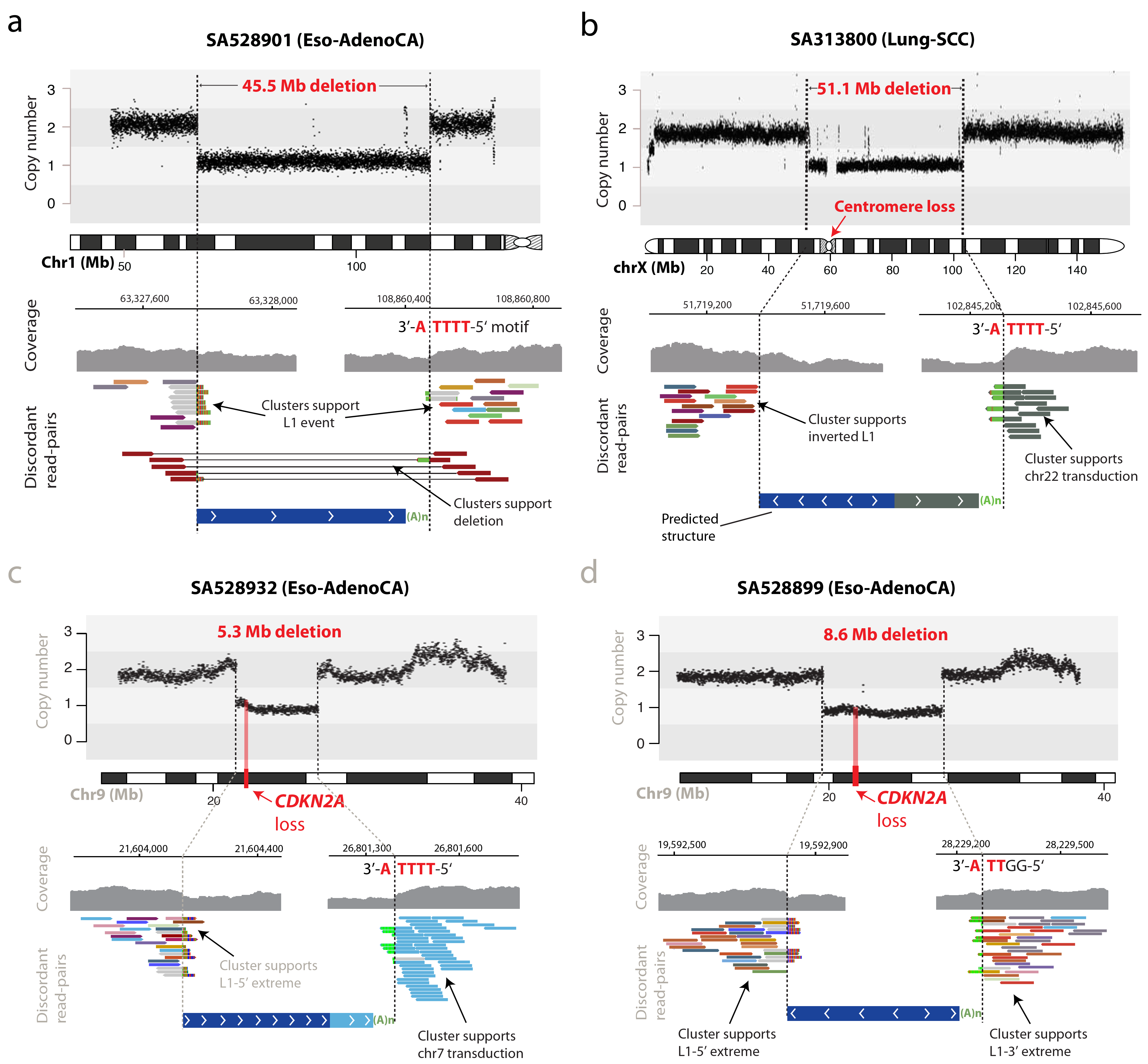
Somatic integration of L1 causes loss of megabase interstitial chromosomal regions in cancer. (a) In an esophageal tumour, SA528901, we find a 45.5 Mb interstitial deletion on chromosome 1 that is generated after the integration of a short L1 event. We observe a pair of clusters of discordant read-pairs whose mates support both extremes of the L1 insertion. Different colours for different reads from the same cluster indicate that mates are mapping a repetitive element; in this case, a L1 retrotransposon. Because the L1 element event is smaller than the library insert size, we also identify read-pairs that span the L1 event and support the deletion. This figure uses default read colours by Integrative Genomics Viewer (igv)^46^, where paired-end reads are coloured by the chromosome on which their mates can be found. See **Supplementary Fig. 2a** for additional information on how to interpret sequencing data. L1-endonuclease 5’-TTTT/A-3’ motif identifies a target-primed reverse transcription (TPRT) L1-integration mechanism. (b) In an esophageal tumour, SA313800, a transduction from chromosome 22 and its companion L1 element is integrated on chromosome X, promoting a 51.1 Mb deletion that removes the centromere. One negative cluster supports a small region transduced from chromosome 22 that bears a poly(A) tract. Reads from this cluster are coloured in green, because their mates map onto chromosome 22 (green is the igv default colour for this chromosome). The L1-endonuclease 5’-TTTT/A-3’ motif was identfied. (c) L1-mediated deletions promote loss of tumour suppressor genes. In esophageal tumour SA528932, the somatic integration at chromosome 9 of a transduction from chromosome 7 and its companion L1 element, promotes a 5.3 Mb deletion involving loss of one copy of the tumour suppressor gene *CDKN2A*. The sequencing data shows a positive cluster of reads whose mates map onto the 5’ extreme of an L1, and a negative cluster that contain split-reads matching a poly(A) and whose mates map onto a region transduced from chromosome 7. Reads from this cluster are coloured in light blue, the default igv colour for mates on chromosome 7. (d) Similarly, in a second esophageal adenocarcinoma, SA528899, the integration of an L1 retrotransposon generates an 8.6 Mb deletion involving the same tumour suppressor gene, *CDKN2A*. The sequencing data reveals two clusters, positive and negative, whose mates support the L1 event integration, together with clipped-reads that precisely mark the insertion breakpoint to base pair resolution.

L1-mediated deletions were, on occasion, driver events, causing loss of tumour suppressor genes. In oesophageal tumour SA528932, the integration of an L1-transduction from chromosome 7p12.3 into the short arm of chromosome 9 caused a 5.3 Mb clonal deletion involving the 9p21.3-9p21.2 region. This led to loss of one copy of a key tumour suppressor gene, *CDKN2A* (Fig. 5c), deleted in many cancer types including oesophageal tumours^34-37^. In this case, the inserted L1 element retained its original structure, meaning that it could have remained active^28^. Interestingly, the sequencing data revealed a somatic transduction arising from this L1 element at its new insertion site, demonstrating that L1 events that promote deletions can be competent for retrotransposition (**Supplementary Fig. 13)**. In a second oesophageal tumour, SA528899, an L1 element integrated into chromosome 9 promoted an 8.6 Mb clonal deletion encompassing the 9p22.1-9p21.1 region that removes one copy of the same tumour suppressor gene, *CDKN2A* (Fig. 5d). Thus, L1-mediated deletions have clear oncogenic potential.

### L1 retrotransposition generates other types of structural variation in human tumours

Somatic retrotransposition can also be involved in mediating or repairing more complex structural variants. In one oesophageal tumour, SA528896, two separate L1-mediated structural variants were present within a complex cluster of rearrangements (Fig. 6a). In the first, an L1 transduction from a source element on chromosome 14q23.1 bridged an unbalanced translocation from chromosome 1p to 5q. A second somatic retrotransposition event bridged from chromosome 5p to an unknown part of the genome, completing a large interstitial copy number loss on chromosome 5 that involves the centromere. This case suggests that retrotransposon transcripts and their reverse transcriptase machinery can mediate breakage and repair of complex dsDNA breaks, spanning two chromosomes.

**Figure 6.**
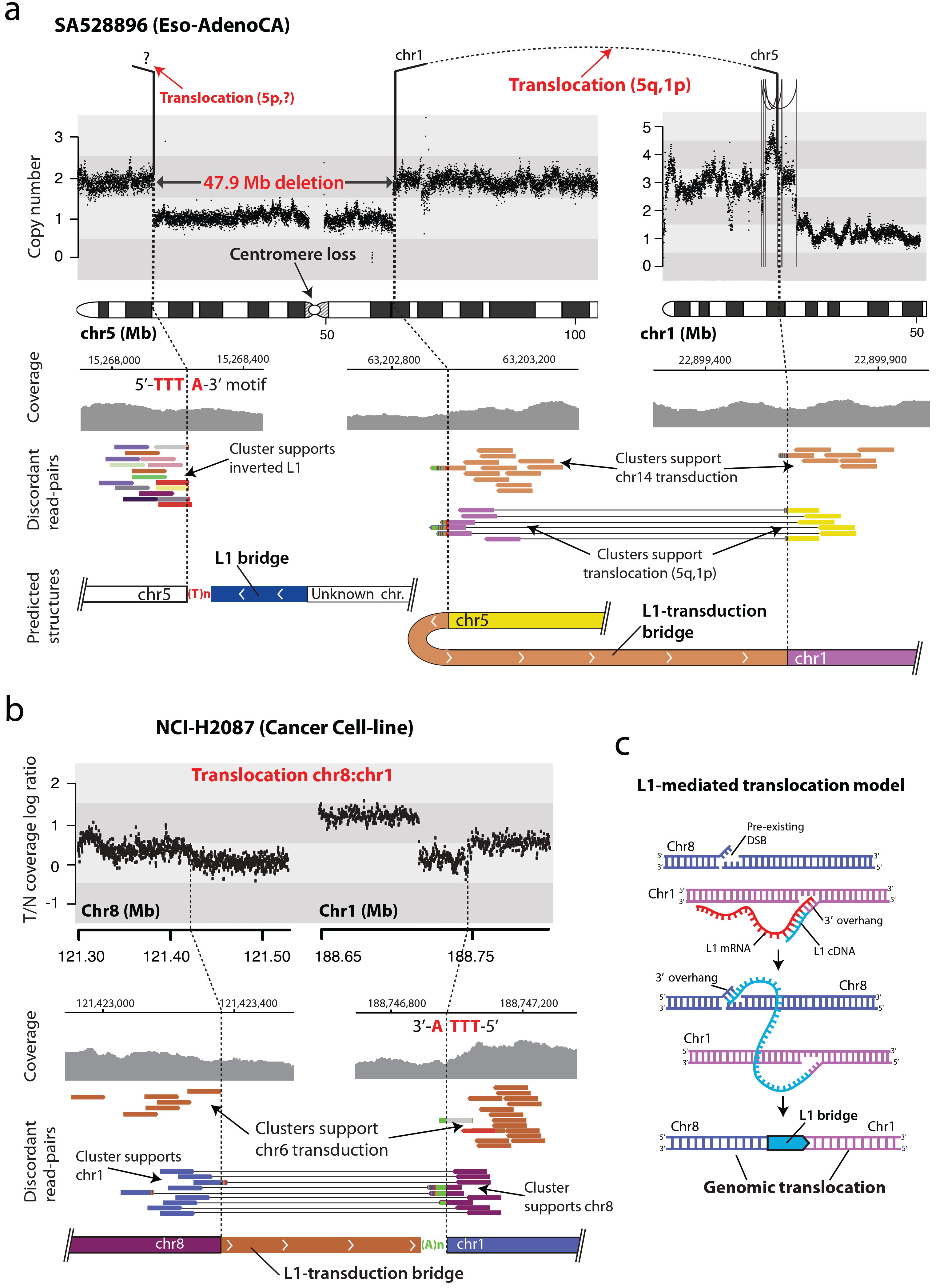
Somatic L1 integration promotes translocations in human cancers. (a) In esophageal adenocarcinoma SA528896, two separate L1 events mediate interchromosomal rearragements. In the first, an L1 transduction from a source element on chromosome 14q23.1 bridged an unbalanced translocation from chromosome 1p to 5q. A second somatic retrotransposition event bridged from chromosome 5p to an unknown part of the genome, completing a 47.9 Mb interstitial copy number loss on chromosome 5 that removes the centromere. This figure uses default read colours by Integrative Genomics Viewer (igv)^46^, where paired-end reads are coloured by the chromosome on which their mates can be found. Different colours for different reads from the same cluster indicate that mates are mapping a repetitive element. See **Supplementary Fig. 2a** for additional information on how to interpret sequencing data. (b) In a cancer cell-line, NCI-H2087, we find an interchromosomal translocation, between chromosomes 8 and 1, mediated by a region transduced from chromosome 6, which acts as a bridge and joins both chromosomes. We observe two read clusters, positive and negative, demarcating the boundaries of the rearrangement, whose mates support the transduction event. In addition, two reciprocal clusters span the insertion breakpoints, supporting the translocation between chromosomes 8 and 1. (c) A model for megabase L1-mediated interchromosomal rearrangements. L1-endonuclease cleavage promotes a 3’-overhang in the negative strand, retrotranscription starts, and the cDNA (-) strand invades a second 3’-overhang from a pre-existing double-strand break on a different chromosome, leading to translocation.

To explore this further, we identified single-L1 clusters with no reciprocal cluster in the cancer cell-lines that were sequenced using mate-pairs with 3 kb and 10 kb inserts. Such events may correspond to hidden genomic translocations, linking two different chromosomes, in which L1 retrotransposition is involved. One of the samples, NCI-H2087, showed translocation breakpoints at 1q31.1 and 8q24.12, both with the hallmarks of L1-mediated deletions, where mate-pair sequencing data identifies an orphan L1-transduction from chromosome 6p24 bridging both chromosomes (Fig. 6b). The configuration has also been confirmed with single-molecule sequencing long reads (**Supplementary Fig. 11**). This interchromosomal rearrangement is likely mediated by an aberrant operation of L1 integration mechanism, where the L1-transduction cDNA is wrongly paired to a second 3´-overhang from a preexisting double strand break generated in a second chromosome^33^ (Fig. 6c).

We also found evidence that L1 integrations can cause duplications of large genomic regions in human cancer. In the oesophageal tumour SA528848 (Fig. 7a), we identified two independent read clusters supporting the integration of a small L1 event, coupled with a coverage drop at both breakpoints. Copy number analysis revealed that the two L1 clusters demarcate the boundaries of a 22.6 Mb duplication that involves the 6q14.3-q21 region, suggesting that the L1 insertion could be the cause of such rearrangement by bridging sister chromatids during or after DNA replication (Fig. 7b). The analysis of the rearrangement data at the breakpoints identified read-pairs that traverse the length of the L1 insertion breakpoints, and the L1-endonuclease motif is the L1 3’ insertion breakpoint, both confirming a single L1 event as the cause of a tandem duplication (Fig. 7a). Interestingly, this duplication increases the copy number of the cyclin C gene, *CCNC*, which is dysregulated in some tumours^38^.

**Figure 7.**
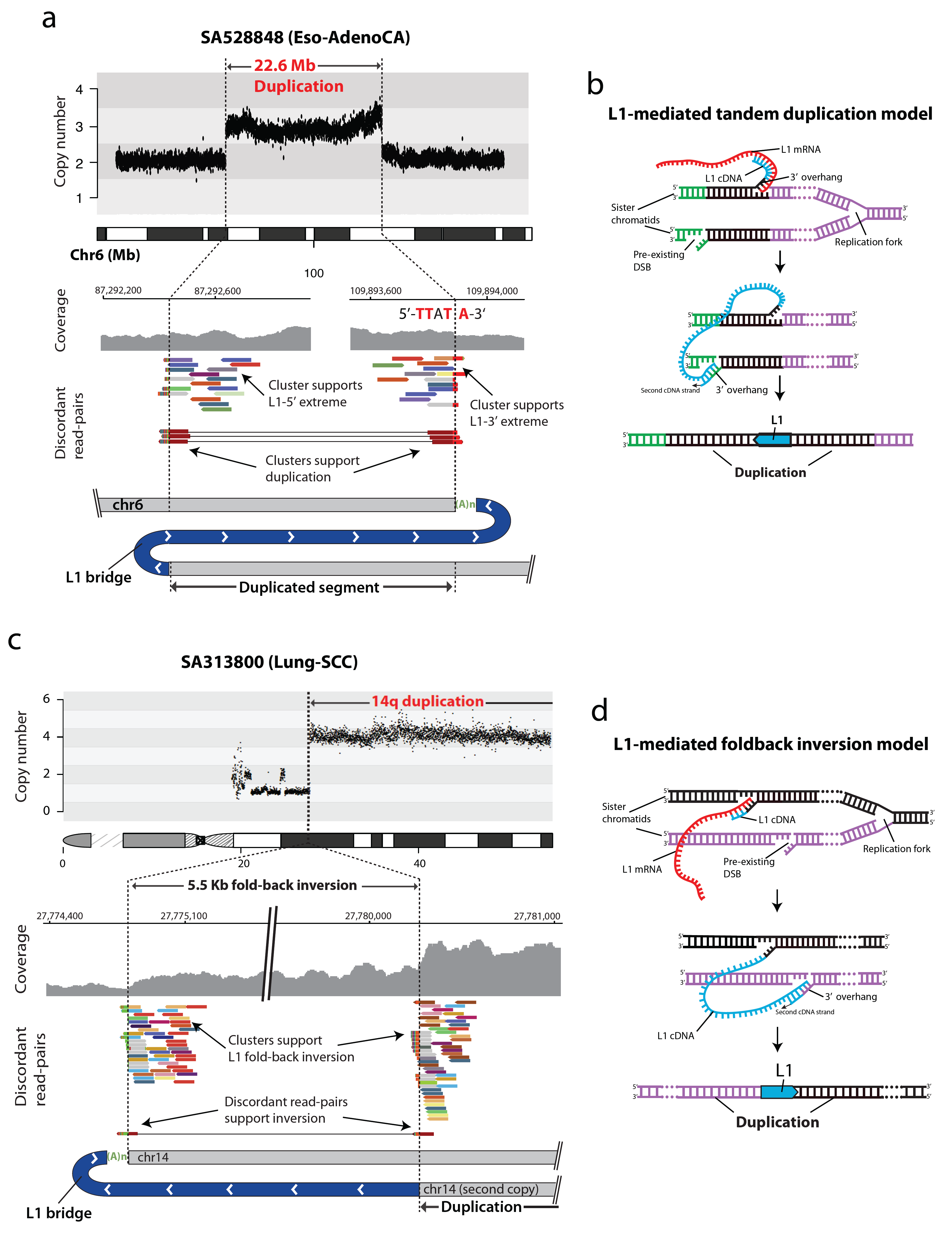
Somatic L1 integration promotes duplications of megabase regions in human cancers. (a) In an esophageal tumour, SA528848, we find a 22.6 Mb tandem duplication at the long arm of chromosome 6. The analysis of the sequencing data at the boundaries of the rearrangement breakpoints reveals two clusters of discordant read-pairs whose mates support the involvement of an L1 event. Because the L1 element is shorter than the library size, we also find two reciprocal clusters that align 22.6 Mb apart on the genome and in opposite orientation, spanning the insertion breakpoints and confirming the tandem duplication. This figure uses default read colours by Integrative Genomics Viewer (igv)^46^, where paired-end reads are coloured by the chromosome on which their mates can be found. Different colours for different reads from the same cluster indicate that mates are mapping a repetitive element. See **Supplementary Fig. 2a** for additional information on how to interpret sequencing data. L1-endonuclease 5’-TTT/A-3’ degenerate motif identifies TPRT L1-integration mechanism. (b) Large direct tandem duplication can be generated if the cDNA (-) strand invades a second 3’-overhang from a pre-existing double-strand break occurring in a sister chromatid, and downstream to the initial integration site locus. (c) In lung tumour SA313800, a small L1 insertion causes a 79.6 Mb duplication of the 14q through the induction of a fold-back inversion rearrangement. The analysis of the sequencing data at the breakpoint revealed two clusters of discordant read-pairs (multi-coloured reads) with the same orientation, aligning close together (5.5 kb apart), and demarcating a copy number change where sequencing density is much greater on the right half of the rearrangement than the left. Both clusters of multicoloured reads support the integration of an L1. The observed patterns suggest that the abnormal chromosome is folded-back on itself leading to duplicated genomic sequences in head-to-head (inverted) orientation, and that a somatic L1 event is bridging the rearrangement. (d) L1-mediated fold-back inversion model.

### L1-mediated rearrangements can induce breakage-fusion-bridge cycles that trigger oncogene amplification

L1 retrotranspositions can also induce genomic instability through triggering breakage-fusion-bridge cycles. This form of genetic instability starts with end-to-end fusion of broken sister chromatids, leading to a dicentric chromosome that forms an anaphase bridge during mitosis. Classically, the end-to-end chromosome fusions are thought to arise from telomere attrition^39-41^. We found, however, that somatic retrotransposition can induce that first inverted rearrangement generating end-to-end fusion of sister chromatids. In lung tumour SA313800 (Fig. 7c), we found a small L1 event inserted on chromosome 14q demarcating a copy number change that involves a 79.6 Mb amplification of the 14q arm. Analysis of the sequencing data at the breakpoint revealed two discordant read-clusters with the same orientation, which are 5.5 kb apart and support the integration of an L1. Both discordant clusters demarcate an increment of the sequencing coverage, where density is much greater on the right cluster. The only genomic structure that can explain this pattern is a fold-back inversion in which the two sister chromatids are bridged by an L1 retrotransposition in head-to-head (inverted) orientation (Fig. 7d).

In the previous example (Fig. 7c-d), no further breaks occurred, and the L1 retrotransposition generated an isochromosome (14q). Beyond this, we found examples in which the fusion of two chromatids by an L1 bridge induced further cycles of breakage-fusion-bridge repair. In the oesophageal tumour SA528848, we identified a cluster of reads at the long arm of chromosome 11 with the typical hallmarks of an L1-mediated rearrangement (Fig. 8a). Copy number data analysis showed that this L1 insertion point demarcated a 53 Mb deletion, involving telomeric region loss, from a region of massive amplification on chromosome 11. The amplified region of chromosome 11 contains the *CCND1* oncogene, which is amplified in many human cancers^42^. The other end of this amplification was bound by a conventional fold-back inversion rearrangement (Fig. 8a), indicative of breakage-fusion-bridge repair^43,44^.

**Fig. 8.**
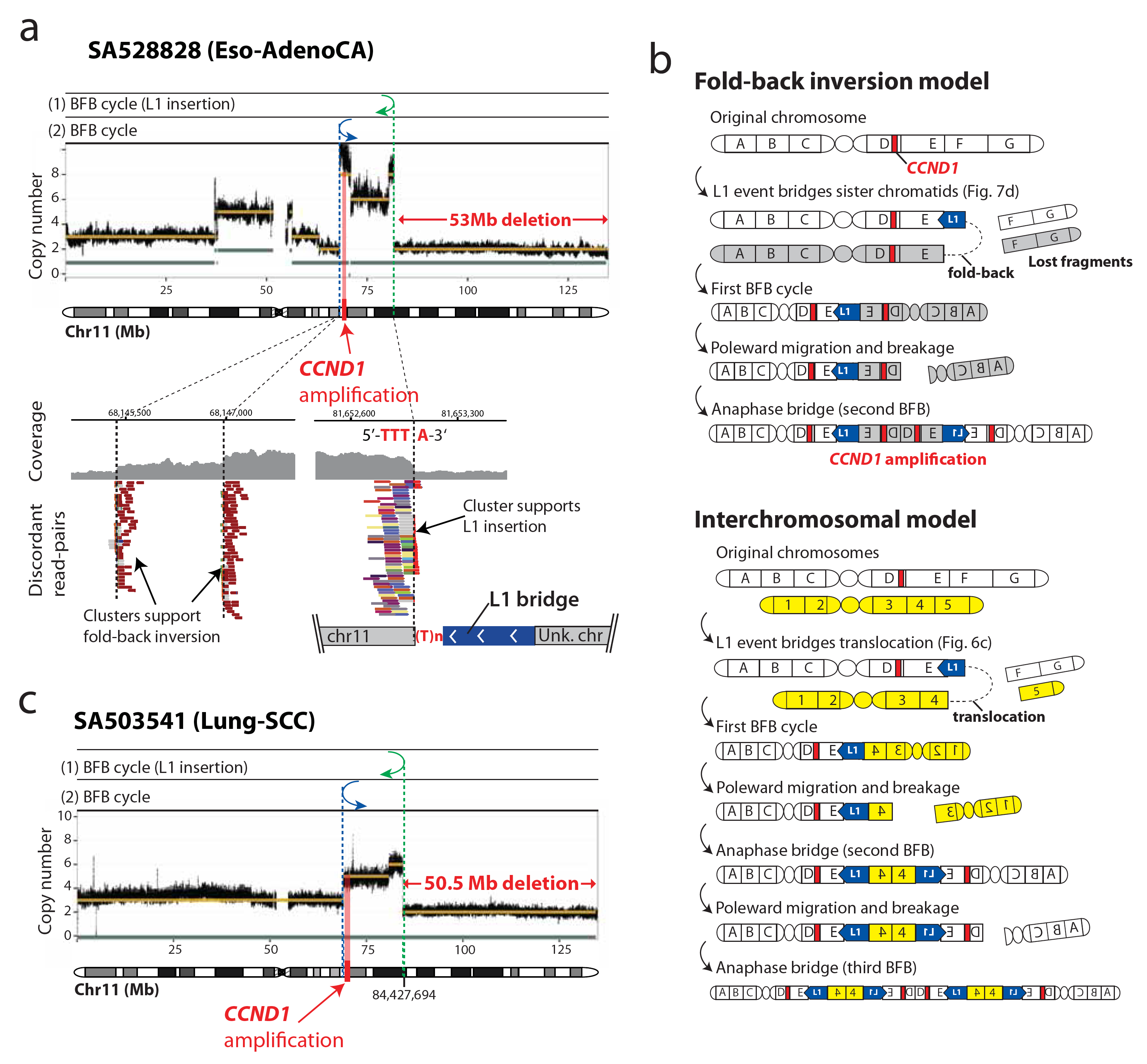
Somatic integration of L1 can trigger breakage-fusion-bridge cycles that lead to oncogene amplification. (a) In an esophageal adenocarcinoma, SA528848, the integration of an L1 retrotransposon on the long arm of chromosome 11 promotes the loss of 53 Mb region that includes the telomere, and activates breakage-fusion-bridge repair leading to amplification of the *CCND1* oncogene. The presence of an L1-endonuclease cleavage site motif 5’-TTT/A-3’ together with a single cluster of discordant reads (positive cluster of multicoloured reads) support the integration of an L1 event. This somatic retrotransposition demarcates a 53 Mb deleted region, involving loss of the telomere, from a region of massive amplification that involves *CCND1*. Around fourteen megabases upstream to the breakpoint of the deletion, we observe the presence of two clusters of read-pairs (i.e., clusters of orange reads) aligning close together and in the same orientation, that demarcate a copy number change; this is a distinctive pattern of a fold-back inversion^43,44^, a rearrangement typically found associated with breakage-fusion-bridge repair. In this fold-back inversion, the coverage show much greater density on the right half of the rearrangement than the left, indicating the abnormal chromosome is folded-back on itself leading to duplicated genomic sequences in head-to-head (inverted) orientation. The patterns described suggest two independent breakage-fusion-bridge cycles, marked with (1) and (2). This figure uses default read colours by Integrative Genomics Viewer (igv)^46^, where paired-end reads are coloured by the chromosome on which their mates can be found. Different colours for different reads from the same cluster indicate that mates are mapping a repetitive element. See Supplementary Fig. 2a for additional information on how to interpret sequencing data. The PCAWG-11 consensus total copy number and the copy number of the minor allele are plotted as gold and gray bands, respectively. (b) Models for the patterns described in a. The fold-back inversion model involves two breakage-fusion-bridge cycles, one induced by L1-mediated fold-back inversion (see Fig. 7d), and a second is by standard breakage-fusion-bridge repair. The intechromosomal rearrangement model involves an interchromosomal rearrangement mediated by an L1, followed by one extra cycle of breakage-fusion-bridge repair. (c) In a lung cancer, SA503541, the integration of an L1 retrotransposon is associated with loss of 50 Mb of the long arm of chromosome 11 that includes the telomere, and activates breakage-fusion-bridge repair leading to amplification of *CCND1*.

These patterns suggest the following sequence of events. During or soon after S phase, a somatic L1 retrotransposition bridges across sister chromatids in inverted orientation, breaking off the telomeric ends of 11q, which are then lost to the clone during the subsequent cell division (‘fold-back inversion model’, Fig. 8b). The chromatids bridged by the L1 insertion now make a dicentric chromosome. During mitosis, the two centromeres are pulled to opposite poles of the dividing cell, creating an anaphase bridge, which is resolved by further dsDNA breakage. This induces a second cycle of breakage-fusion-bridge repair, albeit not one mediated by L1 retrotransposition. These cycles lead to rapid-fire amplification of the *CCND1* oncogene. Alternatively, an interchromosomal rearrangement mediated by L1 retrotransposition (‘interchromosomal rearrangement model’, Fig. 8b) followed by two cycles of breakage-fusion-bridge repair could generate similar copy number patterns with telomere loss and amplification of *CCND1*.

Our data show that L1-mediated retrotransposition is an alternative mechanism of creating the first dicentric chromosome that seeds subsequent rounds of chromosomal breakage and repair. If this occurs near an oncogene, the resulting amplification can provide a powerful selective advantage to the clone. We looked in the PCAWG dataset for other rearrangements demarcating copy number amplification from telomeric deletions that were mediated by L1 integration. We found four more such events across three cancer samples (**Supplementary Fig. 14**). In a lung tumour, SA503541, we found almost identical rearrangements to the one described above (Fig. 8c). Here, a somatic L1 event also generated telomere loss that seeded a second cycle of breakage-fusion-bridge repair. The megabase-size amplification of chromosomal regions also targeted the *CCND1* oncogene, with boundaries demarcated by the L1 insertion breakpoint and a fold-back inversion indicating breakage-fusion-bridge repair. The independent occurrence of these patterns, involving the amplification of *CCND1*, in two different tumour samples, SA528848 and SA503541, demonstrates a mutational mechanism mediated by L1 retrotransposition, which likely contributes to the development of human cancer.

## DISCUSSION

Here we characterize the patterns and mechanisms of cancer retrotransposition on an unprecedented multidimensional scale, across 2,954 cancer genomes, integrated with rearrangement, transcriptomic, and copy number data. This provides new perspective on a long-standing question: Is activation of retrotransposons relevant in human oncogenesis? Our findings demonstrate that major restructuring of cancer genomes can sometimes emerge from aberrant L1 retrotransposition events in tumours with high retrotransposition rates, particularly in oesophageal, lung, and head-and-neck cancers. L1-mediated deletions can promote the loss of megabase-scale regions of a chromosome that may involve centromeres and telomeres. It is likely that the majority of such genomic rearrangements would be harmful for a cancer clone.

However, occasionally, L1-mediated deletions may promote cancer-driving rearrangements that involve loss of tumour suppressor genes and/or amplification of oncogenes, representing another mechanism by which cancer clones acquire new mutations that help them to survive and grow. We expect that structural variants induced by somatic retrotransposition in human cancer are more frequent than we could unambiguously characterise here, given fragment size constraints of the paired-end sequencing libraries. Long-read sequencing technologies should be able to give a more complete picture of how frequent such events are. Relatively few germline L1 loci in a given tumour, typically 1-3 copies, are responsible for such dramatic structural remodeling. Given the role these L1 copies may play in some cancer types, this work underscores the importance of characterizing cancer genomes for patterns of L1 retrotransposition.

## SUPPLEMENTARY TABLES LEGENDS

**Supplementary Table 1.** Description of samples analyzed from the PCAWG dataset

**Supplementary Table 2.** Annotation of somatic retrotransposition events identified in the PCAWG dataset

**Supplementary Table 3.** Counts of somatic retrotransposition events per sample and tumour type in the PCAWG dataset

**Supplementary Table 4.** Distribution of L1 retrotransposition rate and genome organization features in PCAWG genomes

**Supplementary Table 5.** List of germline L1 source elements with transduction counts per tumour sample in the PCAWG dataset

**Supplementary Table 6.** Features of somatic L1-mediated rearrangements identified in the PCAWG cohort

**Supplementary Table 7.** PCR and and single-molecule sequencing validation of 20 L1-meadiated rearrangements in cancer cell-lines

## SUPPLEMENTARY FIGURE LEGENDS

**Supplementary Figure 1. Whole-genome sequencing coverage in tumours and matched-normal genomes in the PCAWG cohort.** (a) Violin plot for the distribution of the mean coverage from all PCAWG tumours analyzed in this study shows a bimodal distribution with maxima at 38 and 60 reads per bp. (b) Distribution of the mean coverage from PCAWG tumours by cancer type. (c) Violin plot for the distribution of the mean coverage from all PCAWG matched-normal samples analyzed in this study shows a mean coverage of 30 reads per genome. (d) Distribution of the mean coverage from PCAWG matched-normal samples by cancer type.

**Supplementary Figure 2. Distribution of somatic retrotranspositions according to subfamily.** (a) L1 subfamilies. Ta-1 and Ta-0 elements – the youngest subfamilies of L1 retrotransposons – represent 97.5% of all L1 somatic mobilizations that were characterized to subfamily level, although we also find 367 L1 events bearing the diagnostic hallmarks of pre-Ta elements, which have been shown to retain retrotransposition activity in modern humans^5^. Elements of the lineages Alu (mainly subfamily AluYa5) and SVA (mainly subfamily SVA_F), The category “Ta-?” contains those L1-Ta events for which it was not possible to detect the Ta-0 or Ta-1 diagnostic nucleotides. (b) Alu subfamilies. (c) SVA subfamilies.

**Supplementary Figure 3. Overview of TraFiC-mem.** (a) TraFiC-mem analyzes Illumina paired-end whole-genome sequencing data, aligned with BWA-mem, from a pair of tumour and matched-normal samples, to identify somatically acquired mobile element insertions (MEIs). See On-line methods for a full description of the algorithm. In this figure, and in the remaining figures where sequencing data is shown, we use default read colours by Integrative Genomics Viewer (igv)^46^, where paired-end reads are coloured by the chromosome on which their mates can be found. Thus, retrotransposon-specific clusters (i.e., clusters of reads supporting the integration of a retrotransposon) are conformed of multicoloured reads, because mates can map ambiguously elsewhere in the reference genome where a retrotransposon of the same family is present. i) Identification of candidate somatic MEIs by TraFiC-mem: For illustrative purposes, the patterns of three different representative insertion types are shown: the integration of a Solo-retrotransposon is detected by the identification of two reciprocal clusters (positive and negative) of interchromosomal (multicoloured) reads whose mates map onto retrotransposon of the same type located elsewhere in the genome; the integration of a partnered transduction from chromosome 7 is detected by the identification of two different types of reciprocal clusters: one cluster of multicoloured interchromosomal reads whose mates map onto L1 retrotransposons of the same family elsewhere in the genome, and one single-coloured cluster of reads whose mates are clusterered at a unique region adjacent to a donor source L1 element – in this illustrative example, this last cluster identifies a transduction from chromosome 7, and reads from this cluster are homogeneously coloured in light-blue because that is the default colour in igv for chromosome 7 –; the integration of an orphan transduction from chromosome 7 is detected by the identification of two reciprocal clusters of the same type, but this time both clusters are single-coloured because mates are clustered on the same chromosome. (b) ii) MEI breakpoint analysis: After preliminary detection of putative MEIs, TraFiC-mem seeks for two additional clusters (5’ brakpoint cluster and 3’ breakpoint cluster) of clipped reads in the candidate insertion region, in order to reveal the 5’ and 3’ insertion breakpoint coordinates to base-pair resolution. iii) MEI structural features annotation: some MEI features are obtained, including structural condition (full, partial, inverted), insertion length, orientation, and target-site duplication (TSD) or target-site deletion. iv) Subfamily assignment: The subfamily of L1 insertions is infered through the identification of diagnostic nucleotides for different L1 subfamilies. Alu and SVA subfamilies identification approach uses de novo assembly of insertion supporting reads followed by RepeatMasker. v) MEI locus annotation: The target genomic region is annotated and MEIs inserted within cancer genes, according to the COSMIC database, are flagged. VCF output: A standard VCF file containing MEI calls is produced. Each MEI is associated with valuable pieces information, as their family, subfamily, length, structure, DNA strand, TSD or deletion length, gene annotation, the number of reads supporting the event and the consensus sequences spanning the breakpoint junctions.

**Supplementary Figure 4. Validation and evaluation of TraFiC-mem.** We validated TraFiC-mem calls on a cancer cell-line with high retrotransposition rate (NCI-H2087) and its matched-normal cell-line (NCI-BL2087), which were sequenced using single-molecule whole-genome sequencing with a MinION (Oxford Nanopore Technologies) to a final coverage of ∼9x (average read size ∼11 kb) and ∼8x (average read size ∼4.5), respectively (see On-line methods). Then, TraFiC-mem algorithm was further evaluated by means of in-silico simulations (a) Retrotransposition breakpoint validation approach using Nanopore long-reads. Illustrative example of a Solo-L1 insertion in cancer cell-line NCI-H2087 detected with Nanopore long-reads. Top, TraFiC-mem relies on the identification of discordant read-pairs (DPs) and clipped reads (CRs) to detect an Solo-L1 insertion using Illumina paired-end sequencing data. Bottom, indel reads (IRs) and clipped reads confirmation using Nanopore sequencing data of the same solo-L1 insertion shown above. (b) Overview of the results of the validation approach of TraFiC-mem calls in cell-line NCI-H2087 using Nanopore whole-genome sequencing data. For each type of insertion (solo-L1, partnered transductions, orphan transductions, Alu), the proportion of events that are supported by Nanopore-reads is represented (from zero to more than 5 reads). Retrotransposition events supported by at least one long read and absent in the matched-normal sample were considered true positive (i.e., somatic), while those not supported by Nanopore-reads and/or present in the matched-normal sample were considered false positive. Overall, false discovery rate (FDR) was estimated to be <5% (see on-line methods). The total number of events assessed for each retrotransposition category is shown on the right side of the panel. (c) Precision and recall of TraFiC-mem after in-silico simulation of 10,000 L1 insertions (Solo-L1, partnered and orphan transductions) in tumours of different clonalities at 25%, 50%, 75% and 100%. Precision was >99% and recall >94%, which represents an improvement relative to previous version of the pipeline ^7^ (d) We also evaluated the ability of TraFiC-mem to estimate the length of retrotransposon integrations. The plot shows the correlation between the observed and expected length for 8,025 Solo-L1 insertions simulated in-silico. (e) The ability of TraFiC-mem to estimate the orientation of retrotransposition integrations was evaluated in-silico using the same simulated data as above. Here it is shown the fraction of true positive Solo-L1 events with a predicted orientation consistent (green), and inconsistent (red), with the expected. Orientation consistency was assessed for four clonality levels (25%, 50%, 75%, 100%).

**Supplementary Figure 5. Rates of somatic retrotransposition across PCAWG tumour types.** Violin plots showing the distribution of the number of retrotranspositions per sample across cancer types, for the six different categories of retrotranspositions that were analyzed (Solo-L1, L1-transductions, L1-mediated deletions, Alu, SVA and Processed pseudogenes). Y-axis is repressented in a logarithmic scale.

**Supplementary Figure 6. TP53 mutation is associated with high rates of L1 retrotransposition and structural variants.** Left panels: Distribution of L1 counts (panel above) and structural variant counts (SV; panel below) of three sample groups according to their *TP53* mutational status: wild-type, monoallelic and biallelic driver mutation. Each data point corresponds to one tumour sample. Groups are compared though Mann–Whitney U, and p-values lower than 0.05 and 0.01 are represented as single and double asterisks, respectively. “NS” stands for Non-Significant. Right panels: box-and-whisker plots showing the distribution of L1 counts (panel above) and SV counts (panel below) across tumour types from samples grouped in two categories: *TP53* wild-type and *TP53*-mutated (monoallelic or biallelic). Within a given tumour type, the two groups (wild-type and mutated) are compared using Mann-Whitney U. P-values are shown only when differences between groups are significant. P-values lower than 0.05 and 0.01 are represented using single and double asterisks, respectively.

**Supplementary Figure 7. Correlation between L1 retrotransposition and structural variation burden.** (a) Heatmap showing the correlation between the number of L1 events, the total number of structural variants (SVs) and the number of 5 different types of SVs per sample: deletions (DEL), duplications (DUP), translocations (TRANS), head-2-head inversions (H2HINV) and tail-2-tail inversions (T2TINV). Correlation was assessed both at Pan-Cancer and tumour type levels. Spearman´s correlation coefficients are shown in a blue (negative) to a red (positive) coloured gradient. P-values lower than 0.05 and 0.01 are represented as single and double asterisks, respectively. (b) Scatter plots showing correlations between the number of L1 events and the total number of SVs per sample at both Pan-Cancer and tumour type levels, for those comparisons that were significant in panel a. Each dot represents one-single tumour sample and it is coloured according to tumour histology. Both axes are displayed on a symlog scale.

**Supplementary Figure 8. Some gene expression effects associated with somatic retrotransposition in PCAWG.** (a) Volcano plot representation of the impact of L1 integration in cancer genes showing the gene expression fold-change (x axis) versus inverted significance (y axis). Red dots indicate significant associations under adjusted p-values < 0.1. This analysis revealed two L1 retrotranspositions where the target cancer gene (*ABL2* and *RB1*) is significantly over-expressed compared to the remaining tumours from the same cancer type (Student’s t-test, q < 0.1). (b) Up-regulation of the *ABL2* oncogene in tumour SA494343, a head-and-neck squamous carcinoma (Head-SCC), relative to the expression of the same oncogene in other Head-SCC samples from the PCAWG dataset are also shown. (c) The analysis of RNA-seq data in genes with L1-retrotransposition in promoter regions revealed significant upregulation in additional three genes. (d) We found six instances where bits of somatic retrotranspositions exonize, being incorporated into the host gene transcripts. These include the cancer genes *PTPN11* and *NCOR2*. Green and purple, or green and orange, boxes represent regions involved in the fusion transcript. Thinner green blocks represent 3’ and 5’ UTRs of the host gene. Thin black lines connecting green and blue boxes represent introns. Split and discordant read-pairs supporting a fusion transcript are shown above the representation of the corresponding predicted transcript. (e) We found evidence for expression of 17 processed pseudogenes mobilized somatically, including aberrant fusion transcripts arising from inclusion of 14 processed pseudogenes (shown here) in the target host gene. Arcs with arrows within the circos indicate the processed pseudogene retrotransposition event, connecting the source processed pseudogene (underlined and bold) with the corresponding integration region. Target site is denoted as intergenic when integration occurs out of gene boundaries, or with the host gene name in italics when integration is within a gene. In the outermost layer of the figure, we represent the predicted processed pseudogene-host gene transcripts. For each host gene mRNA, we have inferred the coding potential of each fusion transcript, which is shown underneath the fusion transcript representation. Start codon is denoted as ATG, termination codon as STOP, and uncertain termination is represented using dots.

**Supplementary Figure 9. L1 integration and genomic features.** (a) L1 rate is strongly correlated with L1 endonuclease (EN) motif density (i.e., number of motifs per Mb) distribution across chromosomal domains (Spearman’s ρ = 0.64, P < 0.01). 2D Kernel density estimate (KDE) is displayed over the data points in a blue to red gradient. (b) Correlation between the number of somatic L1 insertions per Mb and replication timing, which is measured through Repli-seq wavelet-smoothed signal (late to early replication) and averaged per Mb. 2D (KDE) is displayed over the data points in a blue to red gradient. (c-e) In each panel, enrichment scores are shown, adjusted for multiple covariates and comparing the L1 insertion rate in bins 1-3 for a particular genomic feature (see genomic features and colour codes in the legend panel above) versus bin 0 of the same feature, which therefore always has log enrichment=0 by definition and is not shown. For replication time, bin 0 is the latest-replicating quarter of the genome. For essentiality, bin 0 is the non-essential genes. For the L1 motif, bin 0 denotes a non-match (4 or more mismatches). MMs stands for the number of mismatches relative to the consensus L1 EN motif. (c) As reported previously ^7^, we confirm fewer L1 events at active promoters (1.63-fold), here detected by the H3K4me3 histone mark 27, yet there is no decrease at active enhancers, marked by H3K27ac; and we detect no significant association between gene essentiality and L1 rates (1.03-fold decrease in essential genes), suggesting that only a minor fraction of the somatic L1 events may be under negative selection. (d-e) Different cancer types and different samples appear remarkably consistent in the biases of L1 events towards later-replicating DNA and towards other epigenomic features examined: (d) Association between L1 insertion rate and multiple genomic features for tumour types. (e) Association between L1 insertion rate and multiple genomic features in samples with at least 100 L1 events. Each data point is coloured according tumour type.

**Supplementary Figure 10. PCR validation of somatic L1-mediated rearrangement calls.** Gel showing PCR results on cancer cell-lines (NCI-H2009 and NCI-H2087) and their matched-normal cell-lines (NCI-BL2009 and NCI-BL2087). We performed validation of 20 L1-mediated rearrangements (for details, see Supplementary Table 7): 16 L1-mediated deletions (Rg1, Rg2, Rg3, Rg4, Rg6, Rg8, Rg9, Rg10, Rg11, Rg13, Rg14, Rg15, Rg16, Rg17, Rg18, Rg19), 1 L1-mediated translocation (Rg20) and 3 independent L1 breakpoints associated with a copy number change from an unknown rearrangement type (Rg5, Rg7, Rg12). For each rearrangement, except those where only one breakpoint is known, at least three regions were amplified in the tumours (see on-line methods): left breakpoint (L), right breakpoint (R), and the target site (T). Arrows are used to highlight the position of some somatic amplicons. Note that the target site could also amplify in the matched-normal sample if the deletion is not too long. “M” denotes the size marker. For illustrative purposes, the oligo design strategy is shown in a panel at the bottom of the figure: amplicons (L, R and T) and oligos – forward (F) and reverese (R) – are represented.

**Supplementary Figure 11. Single-molecule sequencing validation of somatic L1-mediated rearrangement calls.** We sequenced to high-coverage (>1,000x) the PCR amplicons shown in Supplementary Fig. 10 using single-molecule sequencing with a MinION sequencer (Oxoford Nanopore Technologies). We also carried out whole-genome single-molecule sequencing to low coverage of the same two tumour cell-lines (NCI-H2009 and NCI-H2087) subjected to PCR validation. For illustrative purposes, this figure only shows the validation of four representative rearrangements (Rg18, Rg11, Rg13, Rg20). The sequences of the remaining PCR amplicons can be found in Supplementary Table 7. On the left side of each panel, paired-end and Oxford Nanopore reads supporting a given rearrangement are displayed over a virtual reconstruction of the rearrangement breakpoints. On the right side of each panel, nucleotide sequence obtained by single-molecule sequencing validating each event shown in left. Nucleotide colours match those in the virtual reconstruction of the rearrangement (blue for L1, bright-green for poly-A, grey for target region, light-green for transduction). (a) Solo-L1 insertion mediating a 642 bp deletion. (b) Partnered transduction promoting a 2.6 Kb long deletion. (c) A 1.5 Kb deletion generated through an endonuclease independent L1 integration. Long reads confirm the truncation of the L1 element at its 5′ and 3′ ends. (d) Translocation between 1q31.1 and 8q24.12 mediated by an orphan transduction (same rearrangement as in Fig. 6b). Nanopore reads validate the orphan transduction bridge between both chromosomes.

**Supplementary Figure 12. Validation of L1-mediated rearrangements in cancer cell lines by mate-pairs sequencing.** In order to further validate L1-mediated deletions, we performed mate-pair sequencing of long-inserts libraries (3 kb and 10 kb) on two cancer cell-lines with high-retrotransposition rates. For illustrative purposes, here it is shown the validation of a 10.4 kb long deletion promoted by integration of a 768 bp L1 insertion in the cancer cell-line NCI-H2009. The L1 element inserted within the deletion breakpoints is too long to be characterized using standard paired-end sequencing libraries, but the mate-pairs successfully span the breakpoints of the deletion and confirm a single L1 insertion associated with the rearrangement.

**Supplementary Figure 13. Some L1-mediated deletions are transduction-competent.** (a) Circos plot summarizing the three concatenated retrotransposition events shown in the panel b. First event, an L1 transduction mobilized from chromosome 7 is integrated into chromosome 9. Second event, this insertion concomitantly causes a 5.3 Mb deletion in the acceptor chromosome 9. Third event, the L1 element causing the deletion is subsequently able to promote a transduction that integrates into chromosome X. (b) Discordant read-pairs in chromosome 9 supports a 5.3 Mb deletion generated by the integration of a transduction from chromosome 7, and reveals an L1-event with full-length structure. Five kilobases downstream, a positive cluster of reads supports a transduction from this L1-retrotransposition event into chromosome X.

**Supplementary Figure 14. Somatic integration of L1 and telomere loss.** The PCAWG-11 consensus total copy number and the copy number of the minor allele are plotted as gold and gray bands, respectively. (a) In a head-and-neck tumour, SA494271, deletion of 1.9 Mb at the short arm of chromosome 10, which involves the telomeric region, is associated with the somatic integration of an L1 retrotransposon. (b) In another head-and-neck tumour, SA494351, two independent L1 events promote deletion of both ends of chromosome 5. (c) In a Lung squamous carcinoma, SA503541, the aberrant integration of an L1 event bearing 5’ and 3’ transductions causes a complex rearrangement with loss of 50.5 Mb from the long arm of chromosome 11 that includes the telomere. Only the two clusters supporting both extremes of a putative L1-mediated fold-back inversion are shown. Below, a detailed view of the 5’-transduction breakpoint.

## ONLINE METHODS

## PAN-CANCER DATASETS

### Whole genome sequencing dataset

We analysed Illumina whole-genome paired-end sequencing, 100-150 bp, reads from 2,954 tumours and matched normal samples across 31 cancer types ^21^. For the great majority of donors, the tumour specimens consisted of a fresh frozed sample, while the normal specimens consisted of a blood sample. However, in a small number of cases the normal sample originated from tumour-adjacent normal tissue or another non-blood tissue, particularly for blood cancers. Most of the tumour samples came from treatment-free, primary cancers, but there were also a small number of donors with multiple samples of primary, metastatic and/or recurrent tumour. The average coverage was 30 reads per genomes for normal samples, while tumours had a bimodal coverage distribution with maxima at 38 and 60 reads per bp (**Supplementary Fig. 1 and Supplementary Table 1**). BWA-mem v0.7.17 ^47^ algorithm was used to align each tumour and normal sample to human reference build GRCh37, version hs37d5. Based on the robustness of the retrotransposition calls (false discovery rate <5%), we opted to retain all samples preliminarly excluded by the PCAWG Consortium, as they were excluded from single-nucleotide variants and structural variation analyses based on read direction biases from PCR artifacts or poor sequence quality, but were not found to be problematic for retrotransposition analysis. Additional technical details of the sequencing metrics are given in **Supplementary Table 1** and in the PCAWG maker paper ^21^.

### Transcriptome dataset

About half of the donors (1,188) with a whole-genome in PCAWG had at least one tumour specimen with whole transcriptome obtained by RNA-sequencing. Mapping onto the reference (hs37d5) was carried out using two independent read aligners, STAR ^48^ and TopHat2 ^49^. Gene expression was quantified with HTSeq ^50^, and consensus normalized expression values, in fragments per kilobase of transcript per million mapped reads (FPKM), were obtained by averaging the expression from STAR and TopHat2. A more detailed description of RNA-seq data processing is provided by the PCAWG working group 3 (PCAWG-3) ^51^.

### Copy number dataset

We analyzed copy number profiles obtained by the PCAWG working group 11 (PCAWG-11) using a consensus approach combining six different state-of-the-art copy number calling methods ^52^. GC content corrected LogR values were extracted using the Battenberg algorithm ^53^, smoothed using a running median, and transformed into copy number space according to n = (2(1 – *ρ*) + *ψ ρ*)2^*LogR*^ / *ρ*, where *ρ* and *ψ* are the PCAWG-11 consensus tumour purity and ploidy, respectively.

### Structural variants (SVs) dataset

The structural variation call set was generated by the PCAWG working group 6 (PCAWG-6) by merging the SV calls from four independent calling pipelines ^45^. The merged SV calls were further required to have a consistent copy number change.

## ANALYSIS OF SOMATIC RETROTRANSPOSITION

### Detection of mobile element insertions using TraFiC-mem

BAM files from tumour and matched-normal pairs were processed with TraFiC-mem v1.1.0 (https://gitlab.com/mobilegenomes/TraFiC) to identify somatic mobile element insertions (MEIs) including solo-L1, L1-mediated transductions, Alu, SVA and ERV-K, using Illumina paired-end mapping data. In donors where multiple samples of primary, metastatic and/or recurrent tumours were available, each sample was independently processed and a list of non-redundant MEI calls for each donor was generated by merging the MEI calls from multiple samples as follows: those MEIs within a breakpoint offset of ±15 bp were clustered together, and the call supported by the highest number of discordant read-pairs in each cluster was selected as representative in the merging process.

TraFiC-mem starts by identifying candidate somatic MEIs via the analysis of discordant read-pairs. Contrary to previous version of the algorithm ^7^, the new pipeline uses BWA-mem instead of RepeatMasker as search engine for the identification of retrotransposon-like sequences in the sequencing reads. Calls obtained at this step are preliminary, in which MEI features are outlined and insertion coordinates represent ranges surrounding the breakpoints. Then, a new module of TraFiC-mem, called MEIBA (from Mobile Element Insertion Breakpoint Analyzer) (https://github.com/brguez/MEIBA/tree/master/src/python), is used to identify the integration breakpoints to base-pair resolution, and to perform a detailed characterization of MEI features, including structure, subfamily assignment, and insertion site annotation. TraFiC-mem is illustrated in **Supplementary Fig. 3**.

#### (i) Identification of MEI candidates via discordant reads analysis

- Identification of Solo-element events: TraFic-mem identifies reads from BWA-mem mapping that are likely to provide information pertaining to mobile elements site inclusion. Two different read-pair types are considered for the identification of insertions, named INTER_CHROM (each end of the pair are mapped to different chromosomes, where the end with the highest mapping quality (MAPQ) is considered the anchor in the reference genome), and ABERRANT (both ends of the pair are impropertly mapped to the same chromosome, where the end with the highest MAPQ is considered the anchor). In all cases the anchor end must have a MAPQ higher than zero. The pair is excluded if any of the reads is not a primary alignment, fails platform/vendor quality checks, or is PCR or optical duplicate. Then, all non-anchor reads with an anchored mate MAPQ > 0 are interrogated for the existence of mobile element-like sequences. At this step, non-anchor reads are realigned with BWA-mem v0.7.17^7^ into a database containing a set of human full-length mobile element consensus sequences, including L1, Alu, SVA and ERV-K. After BWA-mem search, all anchored reads with mates containing sequences from the same mobile element type are clustered together if (a) they have the same orientation – positive or negative – and (b) the distance relative to the nearest mapped read of the same cluster is equal or less than the average read size. Two main cluster categories are considered, namely positive and negative clusters (i.e., anchor reads mapped onto the reference genome with positive and negative orientation, respectively). Initially, a range of genome coordinates is associated with each single cluster (breakpoint coordinates are refined in a later step – see “Reconstruction of MEI breakpoints via clipped-reads analysis”). Ranges are conformed by a lower coordinate (P_L_POS and N_L_POS, respectively for positive and negative clusters) and an upper coordinate (P_R_POS and N_R_POS, for positive and negative). One positive and one negative cluster are reciprocal if P_R_POS ≥ N_L_POS and abs(N_L_POS – P_R_POS) ≤ 2(average read size). Only positive and negative clusters conformed of 4 or more reads are employed in the assignment of reciprocal clusters. Each reciprocal cluster identifies a candidate mobile element insertion (**Supplementary Fig. 3a**).

- Identification of L1 transductions: Two types of L1-mediated transductions were identified ^7^, namely partnered transductions, in which an L1 and downstream nonrepetitive sequence are retrotransposed together, and orphan transductions, in which only the unique sequence downstream of an active L1 is retrotransposed without the cognate L1. As above, two different read-pair types, INTERCHROM and ABERRANT are considered. In all cases the anchor read and the corresponding mate must have a MAPQ ≥ 37. Then, anchored reads are clustered together if (a) they share the same orientation, (b) the distance relative to the nearest mapped read of the cluster is equal or less than the average read size, and (c) their mates are also clustered together. Two main cluster categories are considered, namely positive and negative clusters, as above. The integration breakpoint of a potential partnered transduction is defined by reciprocal clusters conformed of one-single cluster of (a) INTER_CHROM and/or ABERRANT reads that support an L1 and (b) one-single cluster of INTER_CHROM and/or ABERRANT reads that supports the integration of a unique DNA region from elsewhere in the genome that is located downstream to an L1 source element locus. One L1 cluster and one INTER_CHROM and/or ABERRANT cluster are reciprocal if they are in opposite orientation and P_R_POS ≥ N_L_POS and abs(N_L_POS – P_R_POS) ≤ 2(average read size). Only positive and negative clusters conformed of 4 or more reads are employed in the assignment of reciprocal clusters. Reciprocal clusters represent preliminary transduction calls that must pass the filters described below to be finally selected. The integration breakpoint of a potential orphan transduction is defined by two reciprocal clusters conformed of INTER_CHROM and/or ABERRANT reads. In this case, two clusters of INTER_CHROM and/or ABERRANT reads are reciprocal if (a) they are in opposite orientation, (b) P_R_POS ≥ N_L_POS and abs(N_L_POS – P_R_POS) ≤ 2(average read size), and (c) the two single clusters that constitute their mates are mapped within a distance of 10 kb to each other (**Supplementary Fig. 3a**).

TraFiC-mem performs the actions described above in both, the tumour and the matched normal genomes. In order to remove potential germline calls, MEI candidates are filtered out from the tumour sample if: (a) they are located within 200 bp of a cluster from the same retrotransposon family in the matched normal sample that is supported by at least 3 reads; and/or (b) there is a polymorphic insertion from the same retrotransposon family within a range of 200 bp that is present in ‘TraFiC-ip db’ ^7^, dbRIP ^54^, the 1,000 Genomes Project Phase 3 callset ^55,56^ or the dataset of germline events identified by our group across PCAWG ^22^. Finally, we noticed the existence of mapping artefacts leading to quite frequent false positive insertion calls (particularly in Alu and SVA calls), located within or between repeats of the same family in the reference genome. So, an additional filter is applied to remove those insertions located within a range of ±150 bp of an element from the same family that shows ≥85% of nucleotide identity relative to the consensus sequence of the family.

#### ii) Reconstruction of MEI breakpoints via clipped-reads analysis

A new module of TraFiC-mem, called MEIBA, is used to identify the breakpoints of an insertion to base-pair resolution. The algorithm uses two classes of reads mapped with BWA-mem, namely soft and hard clipped reads, which overlap a putative insertion breakpoint. These reads consist of two segments, one that aligns onto the insertion target region and a second that aligns onto a mobile element elsewhere in the reference genome. Thus, once MEI candidates have been identified, TraFic-mem seeks for two additional clusters of clipped reads (CRs) that would indicate the exact insertion breakpoint coordinates (each individual insertion has two breakpoints, namely 5’ and 3’ breakpoints). Soft and hard CRs are extracted within a range of ±50 bp to the positive cluster P_R_POS cordinate identified in step “i” of the pipeline via discordant read-pairs. Reads marked as duplicates and reads clipped both at the beginning and ending extremes are filtered out, as they usually constitute mapping artefacts. The same approach is applied to the negative cluster N_L_POS coordinate of the same putative MEI, and both sets of reads, belonging to positive and negative clusters, are merged into a non-redundant dataset as follows: CRs are organized into clusters supporting the same breakpoint position using a maximum breakpoint offset of 3 bp. Those clusters in the tumour that are also detected in the matched-normal genome, and/or those clusters with an overrepresentation of CRs overlapping the breakpoint (we used a cut-off of more than 500 CRs), are excluded. Then, for each breakpoint cluster, supporting CRs are submitted to multiple sequence alignment using MUSCLE v3.8.31 ^57^, and a consensus sequence spanning the insertion breakpoints is constructed with “Cons” from the EMBOSS suit v6.6.0 ^58^. The consensus sequences obtained are processed to assess if they span the target genome region and mobile element breakpoint junction (5’ breakpoint), or the target region and poly(A) tail junction (3’ breakpoint). If more than one 5’ and/or 3’ breakpoints are generated, the one supported by the highest number of CRs is selected. Finally, insertion breakpoints are required to be consistetly supported by at least two CRs, and candidate MEIs are filtered out if they do not have at least one of the two insertion breakpoints characterized to base pair resolution.

#### iii) MEI structural features annotation

MEI structural features including insertion length, structure condition (full-length, partial, inverted), orientation, and size of target site duplication (TSD) or target site deletion, are determined for the insertions with both breakpoints successfully reconstructed. In order to compute the insertion size, the consensus sequence spanning the 5’ breakpoint is realigned to the corresponding L1, Alu, SVA or ERV-K reference sequence using Blat v34.0 ^59^. Next, as retrotransposons usually only get truncated at their 5’-extreme, the insertion length is computed as the distance between the beginning of the alignment and the end of the reference sequence. Insertions spanning less than 98% of the consensus sequence for each family of retrotransposons are considered 5’-truncated, and/or if the sequence aligns in opposite orientation than the insertion DNA strand are considered 5’-inverted; otherwise, insertions are catalogued as full-length. MEIs with positive orientation are supported by clusters at the 5’ and 3’ breakpoints whose CRs are clipped at their ending (end-clipped) and their beginning (beg-clipped), respectively; while MEIs with negative orientation show the opposite clipping pattern. TSD and target-site deletion sizes are estimated as the distance between the two insertion breakpoints. Insertions with TSD show a breakpoint coordinate supported by the end-clipped cluster that is higher than the breakpoint coordinate supported by the beg-clipped cluster. Target site deletions show the opposite pattern.

#### iv) MEI subfamily assignment

Two different strategies are applied to infer the subfamily of the inserted L1, Alu, SVA and ERV-K element. For L1 insertions, discordant read-pairs from supporting reciprocal clusters are realigned onto an L1 consensus sequence (GenBank identifier: L19088.1) using BWA-mem v0.7.17. The resulting SAM is converted into a binary sorted BAM file using samtools v1.7 ^60^. Then, genotype likelihoods at each genomic position are computed with samtools mpileup, reference and variant sites are called with bcftools v1.7 consensus caller ^61^ and filtered requesting a quality score higher than 20 and a minimum read depth of 2. Finally, subfamily inference is done based on the identification of subfamily diagnostic nucleotide positions ^62^: L1 integrations bearing the diagnostic “ACG” or “ACA” triplet at 5,929-5,931 position are classified as “pre-Ta” and “Ta”, respectively. Ta elements are subclassified into “Ta-0” or “Ta-1” according to diagnostic bases at 5,535 and 5,538 positions (Ta-0: G and C; Ta-1: T and G). When sequencing reads do not cover the diagnostic nucleotides, subfamily cannot be inferred. For Alu, SVA and ERV-K, discordant read-pairs from reciprocal clusters supporting the insertions are assembled with velvet v1.2.10 ^63^, using a k-mer length of 21 bp, and the resulting contig is processed with RepeatMasker v4.0.7 to determine the subfamily. If multiple RepeatMasker hits are obtained, the one with the highest Smith-Waterman score is selected as representative. MEIs will be discarded if preliminary family assigment in “i” and subfamily assigment are not consistent.

#### v) MEI locus annotation

The target genomic region is annotated using the software ANNOVAR v2016-02-01 ^64^ and GENCODE v19 basic annotation ^65^. MEIs inserted within cancer genes, according to the Cancer Gene Census COSMIC database v77 ^66^, are flagged.

#### TraFiC-mem output

The primary TraFiC-mem output is a standard Variant Call Format (VCF) v4.2 file containing all somatic MEI calls coordiantes with annotation features, including family, subfamily, insertion length, structural condition, orientation, size of TSD or deletion, gene annotation, number of supporting reads, and consensus sequences spanning the breakpoint junctions. Additional information is provided for L1-mediated transductions, which includes the transduced sequence length, the genomic position of the source element, and source element. MEI candidates that were filtered out are also reported together with filtering reasons.

#### TraFiC-mem availability and distribution

TraFiC-mem is implemented using Snakemake ^67^, a flexible Python-based workflow language, that allows to execute the pipeline from single-core workstations to computing clusters, without the need to modify the workflow. In order to enhance reproducible research, TraFiC-mem and its third party dependencies are also distributed as a Docker image (https://hub.docker.com/r/mobilegenomes/trafic). TraFiC-mem is distributed together with complete documentation and tutorial (https://gitlab.com/mobilegenomes/TraFiC).

### Identification of germline and somatic L1 source elements

Because L1-mediated transductions are defined by the retrotransposition of unique, non-repetitive genomic sequence, we can unambigously indentify the L1 source element whence they derive ^7^. The method relies on the detection of unique DNA regions retrotransposed somatically elsewhere in the cancer genome from a single locus matching the 10 kb downstream region of (i) a reference full-length L1 element, or (ii) a putative non-reference polymorphic L1 element detected by TraFiC-mem across the matched normal samples in the PCAWG cohort ^22^. When transduced regions were derived from the downstream region of a putative L1 event present in the tumour genome but not in the matched normal genome, we catalogued these elements as somatic L1 source loci.

### Identification of processed pseudogene insertions

An additional, separate module of TraFiC-mem was implemented for the identification of somatic insertions of processed pseudogenes (PSD). The method relies on the same principle as for the identification of somatic MEI events, via detection of two reciprocal clusters of discordant read-pairs, namely positive and negative, that supports an insertion event in the reference genome. However, the method differs from standard MEI calling in where the read-mates map, as here mates are required to map onto exons belonging to the same source protein-coding gene in GENCODE v19. To avoid misclassification with standard genomic rearrangements that involve coding regions, we use MEIBA – described above – to reconstruct the insertion breakpoint junctions looking for hallmarks of retrotransposition, including the poly(A) tract and duplication of the target site. Candidate insertions without a poly(A) tail are discarded.

### Identification of L1-mediated deletions

Independent read clusters, identified with TraFic-mem, supporting an L1 event (i.e., clusters of discordant read-pairs with no apparent reciprocal cluster within the proximal 500 bp, and whose mates support a somatic L1 retrotransposition event) were interrogated for the presence of an associated copy number change in its proximity. Briefly, we looked for copy number loss calls from PCAWG-11 (see “Copy number dataset” above) for which the following conditions were fulfilled: (i) the upstream breakpoint matches an independent L1 cluster in positive orientation, the corresponding downstream breakpoint, if any, from the same copy number change matches an independent L1 cluster in negative orientation, and (iii) the reconstruction of the structure of the putative insertion causing the deletion is compatible with one-single retrotransposition event. We used MEIBA – described above – to reconstruct the insertion breakpoint junctions in order to confirm the ends of the events and identify hallmarks of retrotransposition, including the poly(A) tract and duplication of the target site.

Further to the strategy described above, an additional strategy was adopted to identify L1-mediated deletions shorter than 100 kb, as follows. Coverage drops in the proximity of each independent cluster were detected by, first, normalizing read depth on each side of the cluster using the matched normal sample as a reference. Then, the ratio between the normalized read depth on both sides of the cluster was computed. This calculation was performed for window sizes ranging between 200 and 5,000 bp, with the windows on both sides of the cluster always having the same length. An immediately adjacent ‘buffer’ region of 300 bp was defined on each side of the cluster, and reads within these regions were omitted in read depth calculations, in order to prevent false positives due to sequence repeats at the cluster location. Subsequently, pairs of independent reciprocal (positive–negative) clusters were selected for which (i) the two clusters were located less than 100 kb apart, (ii) a potential drop in read depth ratio was identified, extending from the positive cluster to the negative cluster (statistical significance of read depth ratios was estimated non-parametrically, as described below), and (iii) the reconstruction of the structure of the putative insertion causing the deletion was compatible with a single L1 event. For each selected cluster pair, the continuity and reliability of the copy number drop was assessed by measuring the normalized read depth ratio between non-overlapping 500 bp windows spanning the region between the positive and negative clusters (i.e. within the putative deletion) and windows located upstream and downstream of the positive and negative cluster (i.e. outside the putative deletion), respectively. The significance of each drop in read depth ratio was estimated non-parametrically using a null distribution of normalized read depth ratios. This distribution was obtained for each tumour sample by randomly sampling 100,000 genomic locations, drawn from the copy number segments with the predominant copy number in that particular sample (If the predominant copy number was 1, then segments with a copy number of 2 were used instead to avoid extreme read depth ratios that could arise from potentially undetected deletions). Specifically, read depth ratios were calculated from this sample of locations by comparing the normalized read depth between two 2,500-bp windows located immediately upstream and downstream of each location. Non-parametric *p*-values were calculated by comparing the observed read depth ratios with the ones in this null distribution, and adjusted via Benjamini–Hochberg (BH) multiple-testing correction. Three groups of output clusters were produced, corresponding to decreasing significance of the candidate deletions: first, pairs of reciprocal clusters where both clusters present an adjusted p-value below 0.1 (‘Tier 1’ candidates); second, pairs of reciprocal clusters where only one cluster presents an adjusted p-value below 0.1 (‘Tier 2’ candidates); and third, any individual clusters presenting an adjusted p-value below 0.1 (‘Tier 3’ candidates). The resulting L1-mediated deletion candidates (Tiers 1 and 2) were subsequently confirmed via visual inspection using the Integrative Genomics Viewer (igv) ^46^.

### Analysis of retrotransposition rate enrichment and depletion across tumour types

For each tumour type with a minimum sample size of 15, we assessed if it is enriched or depleted in retrotransposition compared to the overall retrotransposition burden though zero-inflated negative binomial regression, as implemented in the *zeroinfl* function of the pscl R package. This type of model takes into account the excess of zeros and the overdispersion present in this dataset. The MEI counts per sample were regressed on a binary factor expressing whether they belong to that particular type of cancer or to any other cancer type. On each regression, the magnitude and sign of the z-score indicates effect size and directionality of the association. More specifically, positive z-scores indicate higher number of counts in the samples belonging to a particular cancer type compared to the rest (enrichment), while negative indique a lower number of counts (depletion). Each z-score is accompanied by its p-value to indicate the level of statistical significance.

### Analysis of association between mutation in tumour suppressor genes and retrotransposition and structural variantion rates

In order to assess if the disruption of a particular tumour suppressor gene (TSG) is associated with a high level of retrotransposition, we used the whole-genome panorama of cancer driver events per sample produced by the PCAWG working group 9 (PCAWG-9) ^68^. This panorama, includes both coding a non-coding SNV and INDELS, as well as copy number alterations, structural variants and potentially predisposing germline variants. For each TSG in the Cancer Gene Census database with mutational data, we stratified the samples into two groups, namely mutated-TSG and non-mutated-TSG. Then, we compared the MEI counts distribution between both groups using Mann–Whitney *U* test to identify significant differences. P-values were corrected for multiple testing via BH procedure. Adjusted p-values lower than 0.05 were considered significant. This analysis was done at both individual cancer type and Pan-Cancer levels to identify tumour type specific associations. We further investigated if there is a *TP53* dosage effect as follows: every PCAWG sample was classified into three groups according to *TP53* mutational status, namely wild-type, monoallelic and biallelic driver mutation. Then, MEI counts distribution was compared for all possible group-pair combinations with Mann–Whitney *U.* The same analysis described above was applied to investigate the association between *TP53* mutation and other types of structural variation.

### Analysis of the correlation between L1 insertion and structural variation rate

For each sample, we computed the number of MEIs, the total number of SVs and the number of five different SV classes: deletions (DEL), duplications (DUP), translocations (TRANS), head-2-head inversions (H2HINV) and tail-2-tail inversions (T2TINV), when data was available. Then, the correlation between the number of MEIs and the SV burden was assessed at both individual tumour type and Pan-Cancer levels using Spearman’s rank test.

### Analysis of the association between L1 insertion rate and genomic features

L1 insertion rate was calculated as the total number of somatic L1 insertions, identified across the complete PCAWG cohort, per 1-Mb window. L1 endonuclease motif density was computed as the number of canonical endonuclease motifs, here defined as TTTT|R (where R is A or G) or Y|AAAA (where Y is C or T), per 1-Mb. Bivariate correlations between L1 insertion rate, endonuclease motif density and replication timing were assessed using Spearman’s rank.

To study the association of L1 insertion rate with multiple predictor variables at single-nucleotide resolution we used a statistical framework based on negative binomial regression, as described in detail previously ^25^. This method was adapted herein such that originally the regression adjusted for content of trinucleotides in each genomic bin, while in this case we instead adjusted for the content of the L1 endonuclease motif. More specifically, we stratified the genome into four bins (0-3) by the closeness of match to the canonical L1 motif, here defined as TTTT|R (where R is A or G). The bin 0 contains dissimilar DNA motifs, which have 4 or more (out of 5) mismatches (MMs), encompassing 1149.7 Mb of the genome. Bin 1, 2 and 3 contain genome segments with exactly 3, exactly 2 and at most 1 MM, encompassing 749.4 Mb, 380.2 Mb and 114.1 Mb of the GRCh37 assembly, respectively. The closest match of either of the two DNA strands was considered.

Histone mark data (ChIP-Seq for H3K9me3, H3K4me3, H3K36me3, H3K27ac) and DNase hypersensitivity (DHS) data for the regional analyses was collected from Roadmap Epigenomics Consortium by averaging fold-enrichment signal over 8 cell types (E017, E114, E117, E118, E119, E122, E125 and E127) and processed by stratifying into four genomic bins, as described previously ^25^. For histone marks and DHS, bin 0 are the areas of the genome with below-baseline signal (Roadmap fold-enrichment compared to input < 1), while bins 1-3 are approximately equal-sized bins covering the remaining parts of the genome with above-average fold-enrichment score. In particular, DHS bins 1-3 encompass 122.8-123.0 Mb each; for H3K36me3 129.1-136.0 Mb each; for H3K4me3 43.2-43.7 Mb each; for H3K27ac 73.6-75.1 Mb each. RNA-Seq data was also collected from Roadmap and processed as previously ^25^ by averaging over 8 cell types (E071, E096, E114, E117, E118, E119, E122, E127): bin 0 consisted of non-expressed genes (FPKM=0) and intergenic DNA that was not explicitly listed as expressed (total 1076.6 Mb), while bin 1 (up to 0.59 FPKM), 2 (up to 5.68 FPKM) and 3 (above 5.68 FPKM) spanned 389.9, 462.1 and 473.8 Mb of the genome, respectively. Replication time (RT) data was processed similarly as histone marks, but collected from ENCODE and processed by averaging the wavelet-smoothed signal over 8 cell types (HeLa S3, HEP G2, HUVEC, NHEK, BJ, IMR-90, MCF-7 and SK-N-SH) and then dividing into four equal-sized genomic bins (quartiles), where bin 0 is the latest-replicating and bin 3 is the earliest replicating. Essential genes were determined by CERES score based on CRISPR essentiality screens, ordering by median score across all 342 cell lines tested ^69^ and then stratifying genes into equal-frequency bins, from less negative to more negative median CERES score (implying commonly essential genes). For the purposes of finding L1 rates in CERES essential genes an additional 1 kb flanking the transcript was also considered together with the gene. All enrichment scores shown in plots compare bins 1-3 for a particular feature (RT, histone marks, gene expression, L1 motif) versus bin 0 of the same feature, which therefore always has log enrichment=0 by definition and is not shown on enrichment plots. The regional analyses are restricted to parts of the genome with perfect mappability scores, according to the CRG Alignability 75 track of the UCSC browser.

### Analysis of the impact of retrotransposition insertions in gene expression

To study the transcriptional impact of a somatic L1 insertion within COSMIC cancer genes and promoters, we used RNA-seq data to compare gene expression levels in samples with and without somatic L1 insertion. For each somatic L1 insertion within a cancer gene or promoter, we compared the gene FPKM between the sample having the insertion (study sample) against the remaining samples in same tumour type (control samples). Using the distribution of gene expression levels in control samples, we calculated the normalized gene expression differences using Student’s t test. For overcoming the problems due to multiple testing, false-discovery rate adjusted p-values (q-values) were calculated through Benjamini–Hochberg, and adjusted p-values <0.1 were considered to be significant.

### Analysis of processed pseudogenes expression

We analysed the PCAWG RNA-seq data to identify and characterize the transcriptional consequences of somatic integrations of processed pseudogenes (PSD). We interrogated RNA-seq split-reads and discordant read-pairs looking for chimeric retrocopies involving PSDs and target genomic regions. For each PSD insertion somatic call, we extracted all the RNA-seq reads (when available) mapping the source gene and the insertion target region, together with the RNA-seq unmapped reads for the corresponding sample. Then, we used these reads as query of BLASTn ^70^ searches against a database containing all isoforms of the source gene described in RefSeq44, together with the genomic sequence in a [-5 kb, +5 kb] range around the PSD integration site. Finally, we looked for RNA-seq discordant read-pairs and/or RNA-seq clipped reads that support the joint expression of processed pseudogene and target site. Only read-pairs with one of the mates aligned into the host gene mRNA with >98% identity were considered. All expression signals were confirmed by visual inspection with Integrative Genomics Viewer.

## VALIDATION OF SOMATIC RETROTRANSPOSITION ALGORITHMS

### In-silico validation of TraFiC-mem

To evaluate precision and recall of our algorithm TraFiC-mem, we reanalyzed a mock cancer genome into which we had previously seeded known somatic retrotransposition events at different levels of tumour clonality ^7^. Briefly, two reference genomes were used: the standard GRCh37 reference genome and an artificial version of it, namely normal and tumour bam files, respectively. To create the artificial, tumoral genome, 10,000 L1 insertion breakpoints – including solo-L1, partnered and orphan transductions – were randomly distributed in the standard reference genome using Bedtools, of which 9,227 were inserted out of un-sequenced gaps. Then, the next-generation sequencing read simulator ART ^71^ was used to generate paired-end read sequencing of both the standard and the artificial reference genomes to a 38x coverage. The simulation FASTQ files were aligned into the standard reference genome with BWA ^72^, resulting in the normal and tumour simulated bam files. As TraFiC-mem requires BWA-mem alignments, we converted the BAM alignments from Tubio et al ^7^ into FASTQ with biobambam v2.0.25 ^73^ and realigned the reads using BWA-mem v0.7.17 with the default configuration. The resulting SAM files were converted into binary sorted BAM files using samtools v1.7 and duplicates were marked with biobambam. Reads from the normal and tumour BAM files were randomly subsampled and merged with samtools at three distinct proportions to also produce tumour samples with 25%, 50% and 75% clonalities. After that, the four possible tumour and matched-normal pairs were processed with TraFiC-mem to call MEIs. For each clonality, the identified MEIs were compared with the list of simulated MEIs to compute the number of true positive (TP), false positive (FP), true negative (TN) and false negative (FN) calls. Finally, the predicted MEI length and orientation were compared with the expected, and precision and recall were computed as follows: Precision = TP / (TP + FP); Recall = TP / (TP + FN).

### Validation of TraFiC-mem calls using single-molecule sequencing

Due to the unavailability of Pan-Cancer DNA specimens, in order to evaluate our algorithm for the identification of retrotransposon integrations, we performed validation of 308 putative somatic retrotranspositions identified with TraFiC-mem in one cancer cell-line (NCI-H2087) with high retrotransposition rate, and absent in its matched normal cell-line (NCI-BL2087) derived from blood, by single-molecule sequencing using Oxford Nanopore technology. Genomic DNA was sheared to 10 kb fragments using Covaris g-TUBEs (Covaris), cleaned with 0.4x Ampure XP Beads (Beckman Coulter Inc). After end-repairing and dA-tailing using the NEBNext End Repair/dA-tailing module (NEB), whole-genome libraries were constructed with the Oxford Nanopore Sequencing 1D ligation library prep kit (SQK-LSK108, Oxford Nanopore Technologies Ltd). We obtained four and five libraries for NCI-H2087 and NCI-BL2087, respectively. Genomic libraries were loaded on MinION R9.4 flowcells (FLO-MIN106, Oxford Nanopore Technologies Ltd), and sequencing runs were controlled using the Oxford Nanopore MinKNOW software v18.01.6. We used the Oxford Nanopore basecaller v2.0.1 to identify DNA sequences directly from raw data and generate fatsq files. Files with quality score values below 7 were excluded at this point. Minion adapter sequences were trimmed using Porechop v0.2.3 (https://github.com/rrwick/Porechop). Then, we used minimap2 v2.10-r764-dirty^74^ to map sequencing reads onto the hs37d5 human reference genome, and the SAM files were converted to BAM format, sorted and indexed with Samtools v1.7 for each one of sequencing runs. BAM files were merged, sorted and indexed. After this process, sequencing coverage were 8.2x (NCI-BL2087) and 9.17X (NCI-H2087), and average read size of mapped reads were ∼4.5 kb (NCI-BL2087) and ∼11 kb (NCI-H2087).

Once having the whole-genome BAM files, for each one of the 308 putative somatic retrotransposition call identified with TraFic-mem, we interrogated the long-read tumour BAM file to seek for reads validating the event. Two types of MEI supporting clusters of sequencing reads were catalogued (**Supplementary Fig. 4a**), namely (i) “indel-read clusters”, composed of Nanopore-reads completely spanning the insertion, so they can be identified as a standard insertion on the reference, and (ii) “clipped-read clusters”, composed of Nanopore-reads spanning only one of the inserted element extremes, so they get clipped during the alignment in the reference. We observe that short MEI insertions are predominantly supported by indel-read clusters, while longer MEI insertions are mainly supported by clipped-read clusters. MEIs supported by at least one Nanopore-read in the tumour and absent in the matched-normal sample were considered true positive (TP) somatic events, while MEIs not supported by long-reads in the tumour and/or present in the matched-normal were considered false positive (FP) calls. Overall, we find 4.22% (13/308) false positive events, which showed to be particularly frequent in regions with low sequencing coverage. However, we cannot not rule out the possibility that these are true positive events, as they were not found in the matched-normal sample. False discovery rate (FDR) was estimated as follows: FDR = FP / (TP + FP).

### Validation of L1-mediated rearrangements with PCR and single-molecule sequencing

Due to the unavailability of Pan-Cancer DNA specimens, we performed validation of 20 somatic L1-mediated rearrangements, mostly deletions, identified in two cancer cell-lines with high retrotransposition rates (NCI-H2009 and NCI-H2087). We carried out PCR followed by single-molecule sequencing of amplicons from the two tumour cell-lines and their matched normal samples (NCI-BL2009 and NCI-BL2087), using a Minion sequencer from Oxford Nanopore. PCR primers were designed with Primer3 v0.4.0 ^75^, to amplify three regions from each event (namely, 5’-extreme, 3’-extreme and target site) as follows. For the amplification of the 3’-extreme of the event (the one that contains the poly(A) tract), we designed one forward oligo to hybridize the 3’ extreme of the MEI, and a reverse oligo that hybridizes the DNA downstream. In the case of Solo-L1s, we used an L1Hs specific forward oligo matching the 3’-UTR: 5’-GGGAGATATACCTAATGCTAGATGACAC-3’ ^76^, or an alternative forward oligo that matches other region at the 3’-extreme of the element. For the amplification of the target site in the tumour and matched normal, we designed primers to amplify the DNA sequence between both breakpoints (5’ and 3’) of the rearrangement. For the amplification of the 5’-extreme of a MEI in a tumour, we designed one forward oligo to match the non-repetitive region immediately adjacent to the 5’-extreme of the element, and a reverse oligo that hybridizes the 5’ extreme of the MEI.

Each PCR mixture contains 10ng of DNA, 5pmol of each primer, 5U Taq DNA polymerase (Sigma-Aldrich, catalog number D1806) with 1x Buffer containing MgCl_2_, 0.2mM of each dNTPs, and water to a final volume of 25μl. PCR conditions were as follows: initial denaturation at 95°C for 7 minutes; then 30-35 cycles of 95°C for 30 seconds, 60°C for 30 seconds, 72°C for 45 seconds; and a final extension of 72°C for 7 minutes. In some cases, when amplification was tricky, we used Platinum Taq High-fidelity, with a 94°C denaturation and a 68°C extension.

PCR amplicons were sequenced with single-molecule sequencing using a MinION from Oxford Nanopore. Amplicons were pooled and total DNA was cleaned with 0.4x AMPure XP Beads (Beckman Coulter Inc). After end-repairing and dA-tailing using the NEBNext End Repair/dA-tailing module (NEB), the sequencing library was constructed with the Oxford Nanopore Sequencing 1D ligation library prep kit (SQK-LSK108, Oxford Nanopore Technologies Ltd) and loaded on a MinION R9.4 flowcell (FLO-MIN106, Oxford Nanopore Technologies Ltd). Mapping to human reference genome was performed as described above, with minor modifications.

### Validation of L1-mediated rearrangements using mate-pairs

To further validate and characterize L1-mediated rearrangements we performed 10x mate-pair whole-genome sequencing using libraries with two different insert sizes, 4 kb and 10 kb, which can span the integrated L1 element that caused the deletion, allowing validation of the involvement of L1 in the generation of such rearrangements. Mate-pair reads (100 nucleotides long) were aligned to the human reference build GRCh37, version hs37d5, by using BWA-mem v0.7.17 with default settings, with the exception of the mean insert size that was set to 4 kb and 10 kb, accordingly with the insert size library. Then, for each candidate L1-mediated rearrangement we looked for discordant mate-pair clusters that span the bkps and support the L1-mediated event. Each event was confirmed via visual inspection of BAM files using IGV.

## DATA AVAILABILITY

Accession codes to VCF files containing information relative to all somatic insertions identified in this study will be available before publication together with other datasets generated by other PCAWG working groups.

## ACKNOWLEDGEMENTS

J.M.C.T. is supported by European Research Council (ERC) starting grant (Grant Agreement Number: 716290 SCUBA CANCERS ERC_Stg_2016). This work was supported by the Wellcome Trust grant 09805. B.R.M is supported by a predoctoral fellowship from Xunta de Galicia (Spain). A.L.B. is supported by a predoctoral fellowship from the Spanish Ministry of Economy, Industry and Competitiveness (MINECO). R.B. received funding through the National Institutes of Health (U24CA210978 and R01CA188228). M.G.B received funding through MINECO, AEI, Xunta de Galicia and FEDER (BFU2013-41554-P, BFU2016-78121-P, ED431F 2016/019). N.B. is supported by a My First AIRC grant from the Associazione Italiana Ricerca sul Cancro (n. 17658). K.H.B. is supported by P50GM107632 and R01CA163705. J.D. is a postdoctoral fellow of the Research Foundation – Flanders (FWO) and the European Union’s Horizon 2020 research and innovation program (Marie Sklodowska-Curie Grant Agreement No. 703594-DECODE). P.AW.E. is supported by Cancer Research UK. E.A.L. is supported by K01AG051791. I.M. is supported by Cancer Research UK (C57387/A21777). F.M. is supported by A.I.L. (Associazione Italiana Contro le Leucemie-Linfomi e Mieloma ONLUS) and by S.I.E.S. (Società Italiana di Ematologia Sperimentale). Y.S.J. is supported by a grant of the Korea Health Technology R&D Project through the Korea Health Industry Development Institute (KHIDI), funded by the Ministry of Health & Welfare, Republic of Korea (grant number: HI14C1277). J.O.K. is supported by an ERC Starting Grant. S.M.W. received funding through a SNSF Early Postdoc Mobility fellowship (P2ELP3_155365) and an EMBO Long-Term Fellowship (ALTF 755-2014). J.W. received funding from the Danish Medical Research Council (DFF-4183-00233). P.V.L. is a Winton Group Leader in recognition of the Winton Charitable Foundation’s support towards the establishment of The Francis Crick Institute. This is work is supported by The Francis Crick Institute, which receives its core funding from Cancer Research UK (FC001202), the UK Medical Research Council (FC001202), and the Wellcome Trust (FC001202). H.H.K. is supported by grants from the National Institute of General Medical Sciences (P50GM107632 and 1R01GM099875). F.S. was supported by an ERC Starting Grant “HYPER-INSIGHT”.

